# High-throughput discovery of emerging antifungal resistance in crop pathogens

**DOI:** 10.1101/2025.07.18.665620

**Authors:** Guido Puccetti, Daniel Flückiger, Dominique Edel, Camille Delude, Sabina Moser Tralamazza, Thomas Badet, Gabriel Scalliet, Daniel Croll

## Abstract

The rise of antifungal resistance is a global challenge for both human health and food security, because resistance emergence easily outpaces the antifungal development pipeline. Furthermore, resistance arises often in parallel and through alternative mechanisms creating challenges to predict emergence. In agriculture, where vast areas are sprayed by diverse cocktails, antifungal resistance gains are particularly complex. Despite broad efforts, knowledge of resistance mechanisms is often limited to model genotypes and empirical evidence from the field is lacking. Here, we define and validate a high-throughput pipeline for antifungal resistance discovery informed by emerging resistance gains at continental scale. We analyzed a thousand-genome European diversity panel of the major wheat pathogen *Zymoseptoria tritici* and assessed resistance levels against over 29 fungicides covering all major classes. We optimized high-throughput phenotyping assays to comprehensively capture emerging resistance phenotypes. Pangenome-informed genotyping techniques revealed a total of 2192 genes associated with antifungal resistance. This expands by an order of magnitude the current knowledge and establishes a refined atlas of resistance mutations. We generated mutants to recapitulate several of the discovered resistance factors. Hence, our approach captures in-field resistance gains across Europe for all major fungicide classes and can define exact molecular targets. Broad knowledge of resistance gains will guide more sustainable fungicide development pipelines.

## Main

Fungicide resistance is a global concern for medicine and agriculture ^1–5^. Critical challenges arise as the development of new fungicides is slower than the rate of resistance emergence ^6^. Understanding the molecular mechanisms of fungicide resistance is crucial to inform drug design and to mitigate resistance breakdowns. Resistance mechanisms are typically identified through molecular genetics approaches such as random mutagenesis or reverse genetics ^7–9^. Such approaches focus on single genetic backgrounds and may only poorly capture alternative resistance mechanisms emerging through fungicide resistance selection in the field ^10^. Pathogen species can present variation in the molecular target or the expression of detoxification mechanisms leading to resistance heterogeneity ^11^. Monitoring of pathogen populations and assessing changes in fungicide resistance can identify the rise of alternative resistance in the field ^1,12–14^. However, monitoring needs to capture full genomic information to be effective ^5,11,15^.

Mechanisms underpinning the expression of complex traits such as fungicide resistance can be resolved by genome-wide association studies (GWAS) ^5,16,17^. Resistance mapping identified mutations in the ergosterol biosynthesis gene *Cyp51* to azole fungicide resistance in sugar beet pathogens ^18^. In the wheat pathogen *Zymoseptoria tritici*, both the known target of azole resistance and detoxification mechanisms were mutated in field isolates. Comprehensive mapping of emerging resistance factors by GWAS requires a spatially explicit sampling and functional validation of candidate loci ^19,20^. The power of large pathogen GWAS panels to uncover variation in major traits was recently demonstrated by the establishment of thousand-genome panels for the major wheat pathogen *Z. tritici* ^5,21^. The panels revealed global patterns in pathogen emergence, adaptation to climatic factors as well as complex sets of resistance mutations for demethylation inhibitors (DMI) such as azoles. The GWAS panel constructed from European samples identified a total 65 genes associated with resistance to six DMIs in half-maximum growth assessments. This highlights the need to comprehensively capture emerging resistance variants beyond known target genes for agriculturally important fungicides ^11,22^.

Despite recent efforts, the full spectrum of molecular mechanisms contributing to resistance against major fungicide classes remains poorly understood ^5,23^. A major factor are complex genetic contributions to resistance such as structural variation generated by transposable elements (TEs). Complex mutations were found, for instance, in regulatory regions of a drug exporter gene and similar variants arose convergently in multiple pathogen species ^24–26^. Molecular targets can also undergo copy-number variation in response to fungicide selection such as the emergence of drug target paralogs in *Z. tritici* and the opportunistic human pathogen *Candida albicans* ^11,27,28^. Complex genetic variation underpinning trait variation can be mapped using reference-free genotyping approaches such as k-mers^29^. Azole resistance in *Z. tritici* is in part governed by complex genetic variation, which was only recovered using reference-free genotyping approaches ^29^. Beyond advances in variant discovery and mapping, high-throughput resistance screening of large strain collections is necessary. A rigorous evaluation how such pathogen diversity panels should be constructed and how relevant traits should be assessed remains poorly investigated.

Here, we develop and validate a high-throughput pipeline integrating a diverse pathogen panel for GWAS and a robust discovery of complex genetic variants underpinning fungicide resistance. We determined fungicide sensitivity of comprehensive European diversity panel of the wheat pathogen *Z. tritici* for over 29 fungicides and produced a comprehensive fungicide resistance atlas cataloguing an exhaustive set of genetic factors underpinning resistance breakdowns in a geographic and temporal context. We apply functional genetics approaches to assess the validity of key mutations discovered in the resistance atlas.

## Results

### Establishment of a European diversity panel

Following the collection and azole resistance phenotyping of a European diversity panel for *Z. tritici* established from infected wheat fields ^5^, we utilized the strain collection to establish a comprehensive high-throughput pipeline linking natural variation to the genetic basis of fungicide resistance. The European diversity panel captures a timespan of 15 years and originates from extensive field monitoring efforts. The European continent has long experienced fungicide applications, covers a wide range of climates and fungicide regimes (Table S1). Sampling for the European diversity panel was augmented in areas of intense fungicide application and wheat production ^30^ (Table S2, Figure 1A). We optimized a high-throughput phenotyping assay to profile differential fungicide susceptibility across the panel and evaluated complementary genotyping approaches to conduct GWAS (Figure 1B). Finally, we used molecular genetics assays to validate the impact of mapped resistance mutations on fungal growth.

**Figure 1.**
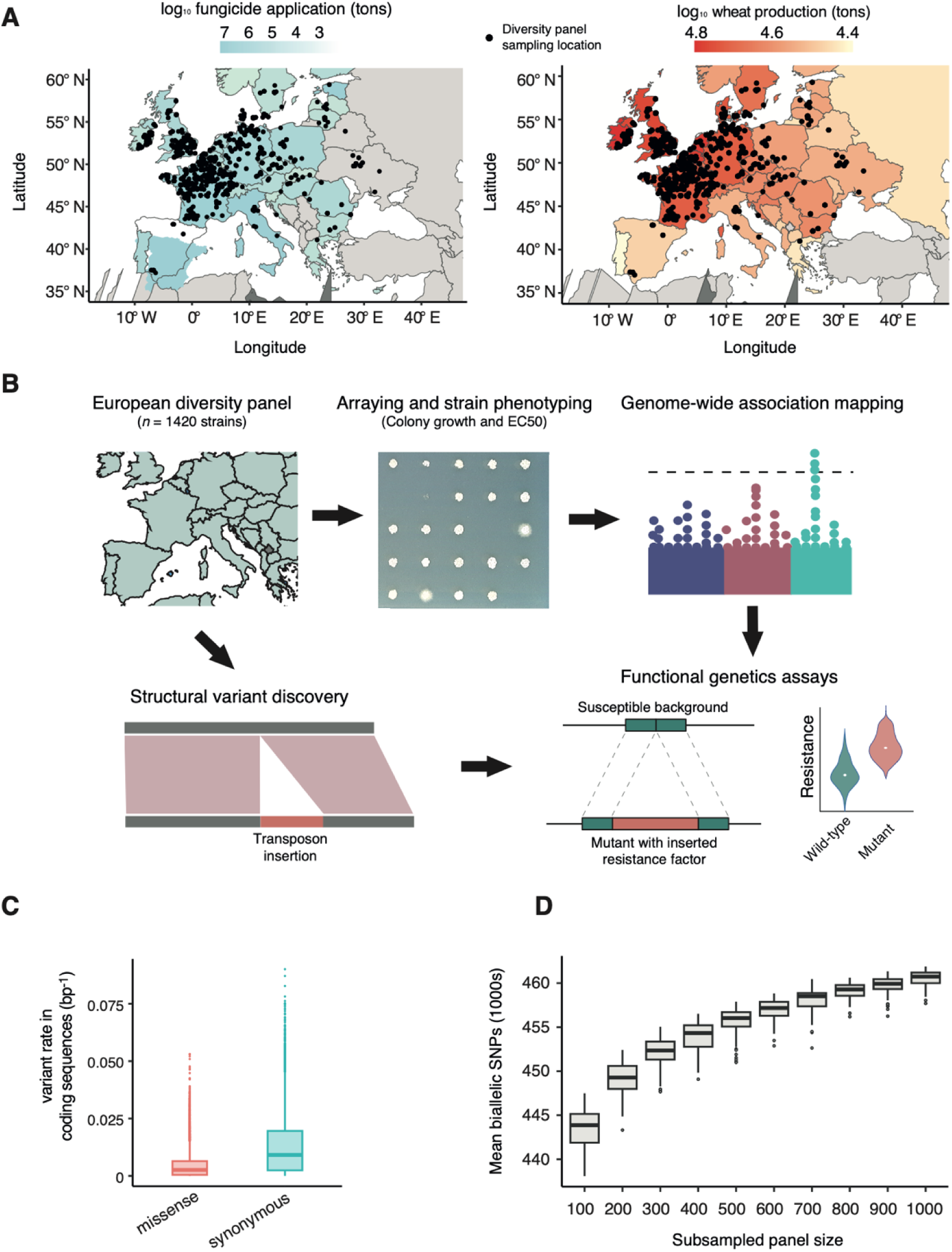
Design of the fungicide resistance discovery pipeline connecting natural variation to the genetic basis of resistance. A: European maps representing European isolate distribution (black dots) and colored the average ton of fungicide applied between 2011 and 2020 and wheat production in between 2017 and 2019. B: Establishing a comprehensive collection of strains across space and time, designing and optimizing a phenotypic assay to profile fungicide resistance, performing GWAS, and conducting functional validation. C: Percentage of synonymous and missense variants, normalized by gene length across the core chromosomes in the European collection. D: Average number of SNP variants derived from a random subsampling of the European collection from 100-1000 strains.

A key factor for successful GWAS applications is the genetic diversity captured by the mapping population ^19^. For the total set of polymorphic SNPs, the diversity panel showed an average of 1.2% synonymous point mutations and 0.4% missense point mutations (Figure 1C). To estimate the impact of strain collection size on the number of segregating variants, we randomly subsampled the European diversity panel to 100-1000 strains. We found an average increase of 0.4% (*n* = 1905) segregating SNPs of the total detected polymorphism for each 100 strains added to the panel. The overall gain between 100 and 1000 strains was of 3.7% (*n* = 17,141) of all polymorphic SNPs (Figure 1D). These results show that even a small but diverse collection of 100 strains sampled across Europe can effectively capture segregating mutations. To further understand what factors maximize variant discovery, we systematically examined the effects of geography and collection timeframe of the sampled strains. We first subset the panel into two timeframes (2005-2015 and 2016-2019) composed of *n* = 370 and *n =* 529 strains, respectively. The latter timeframe covered the first applications of the DMI mefentrifluconazole. Consistent with the experienced exposure in the fields, the 2016-2019 collection showed significantly higher relative growth in mefentrifluconazole compared to the earlier collection (*p* < 0.01; Figure 2A-C). Then, we assessed whether the increased resistance could be associated with the gains of resistance mutations. GWAS analyses identified five SNPs on chromosome 3 in the 2005- 2015 panel, intersecting with a gene encoding a Zinc-finger C2H2-type transcription factor (3_01019). In contrast, the later collection set revealed 152 SNPs associated with mefentrifluconazole resistance located within ∼41 kb of the gene *Cyp51* encoding the target of azoles and an additional three SNPs were identified in the promoter region of the multidrug exporter gene *MFS1* (Figure 2D).

**Figure 2.**
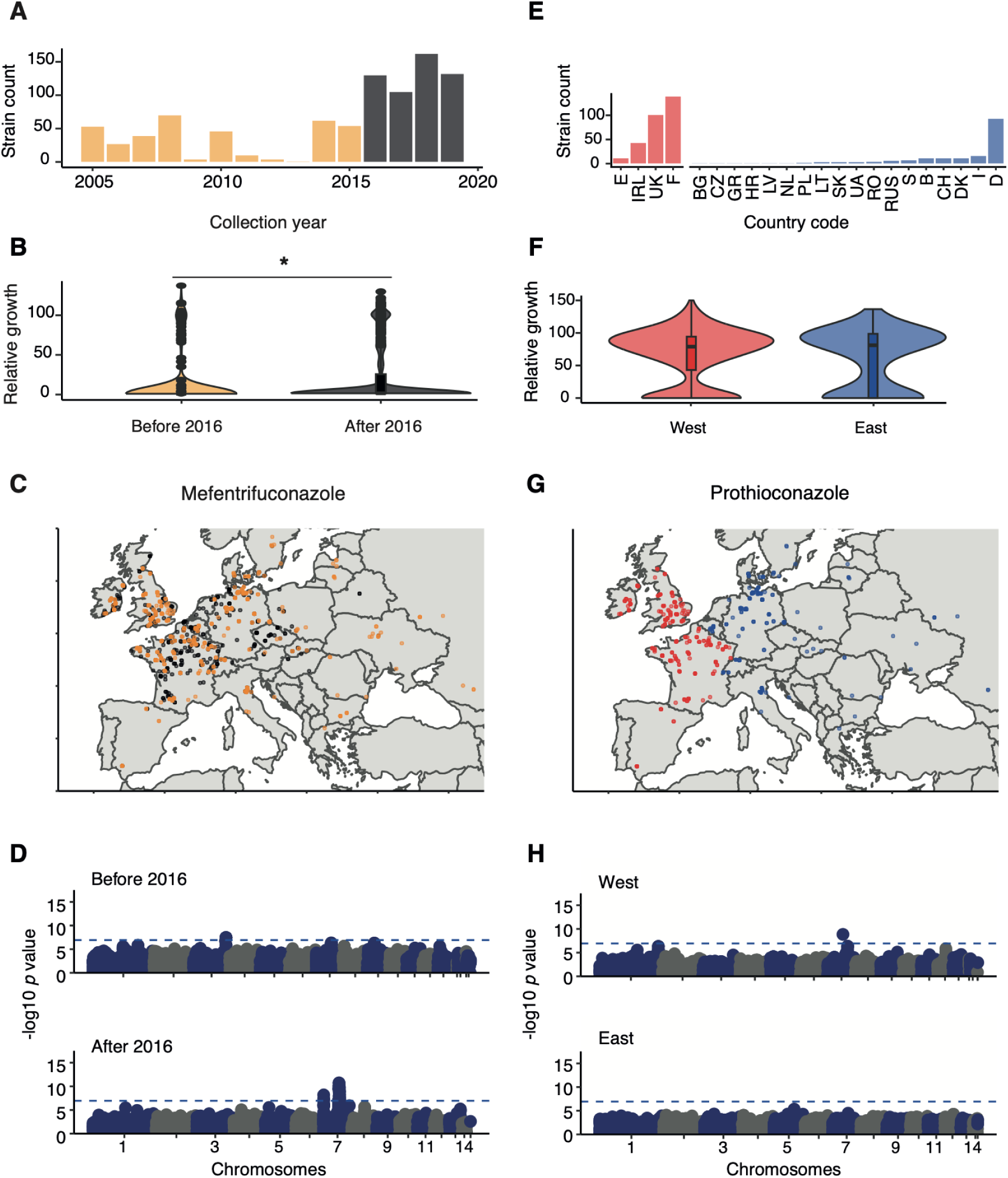
Sampling distribution and association studies of geographically and sampling epoch- constrained panels. (A) Frequency of strains before and after 2015 per year. (B) Relative growth differences for mefentrifluconazole (7 dpi, 0.1 mg.L^-1^) * t-test p-value<0.05. (C) Geographic distribution of strains before and after 2015. (D) Manhattan plot of mefentrifluconazole resistance based on relative growth. For visualization purposes, SNPs have been randomly subsampled. (E) Frequency of strains in the West and East of Europe. (F) Relative growth differences for prothioconazole (7 dpi, 1 mg.L^-1^). (G) Geographic distribution of strains in Western and Eastern Europe. (H) Manhattan plot of prothioconazole resistance based on relative growth values.

Besides the introduction of new fungicides, fungicide application practices vary significantly across Europe, leading to region-specific selection pressure. Hence, local adaptation could produce heterogenous outcomes in adaptation ^31^. In Europe, east-to-west represents a gradient in increased DMI resistance consistent with fungicide application practices ^1^. We compared two sets of strains either collected after 2015 from Western Europe and the British Isles (*n* = 294) or collected in Eastern Europe (*n* = 176) (Figure 2E, F, G). Colony growth on prothioconazole medium showed no significant differences between the two strain sets (*p* = 0.2; Figure 2E). However, GWAS identified an amino acid substitution in the azole target gene *Cyp51* (Ser524Thr) in the Western Europe strain set, while no mutation was significantly associated with resistance in the Eastern European strain set (Figure 2H).

Finally, we examined how the panel size impacts the power of GWAS to detect fungicide resistance mutations. We subset the European diversity panel into sets ranging from 100 to 800 samples, with each subpanel size being randomized 100 times. We performed GWAS for mefentrifluconazole resistance on each panel subset and found that the number of detected resistance SNPs reached a plateau at a panel size of *n =* 700 strains (Figure 3A-B). Mefentrifluconazole resistance mutations converged on three different chromosomal regions, including *Cyp51*, a region ∼35 kb from *Cyp51*, and the *MFS1* gene (Figure 3C). Already at the second smallest subpanel size of *n =* 200 strains, a mean of 6.6 resistance mutations were mapped in *Cyp51* and 4 SNPs were mapped near *MFS1.* For prothioconazole, a panel of *n =* 200 strains was sufficient to identify *Cyp51* resistance mutations (mean of 1.03 SNPs) and a gene of unknown function 12_00177 (mean of 8.6 SNPs) and reaching at plateau in total discovered resistance mutations at *n* = 600 strains (Figure 3D). These findings show that the composition of the GWAS panel significantly affects the ability to identify resistance loci and that panel sizes of at least 800 strains are well-powered for fungicide GWAS.

**Figure 3.**
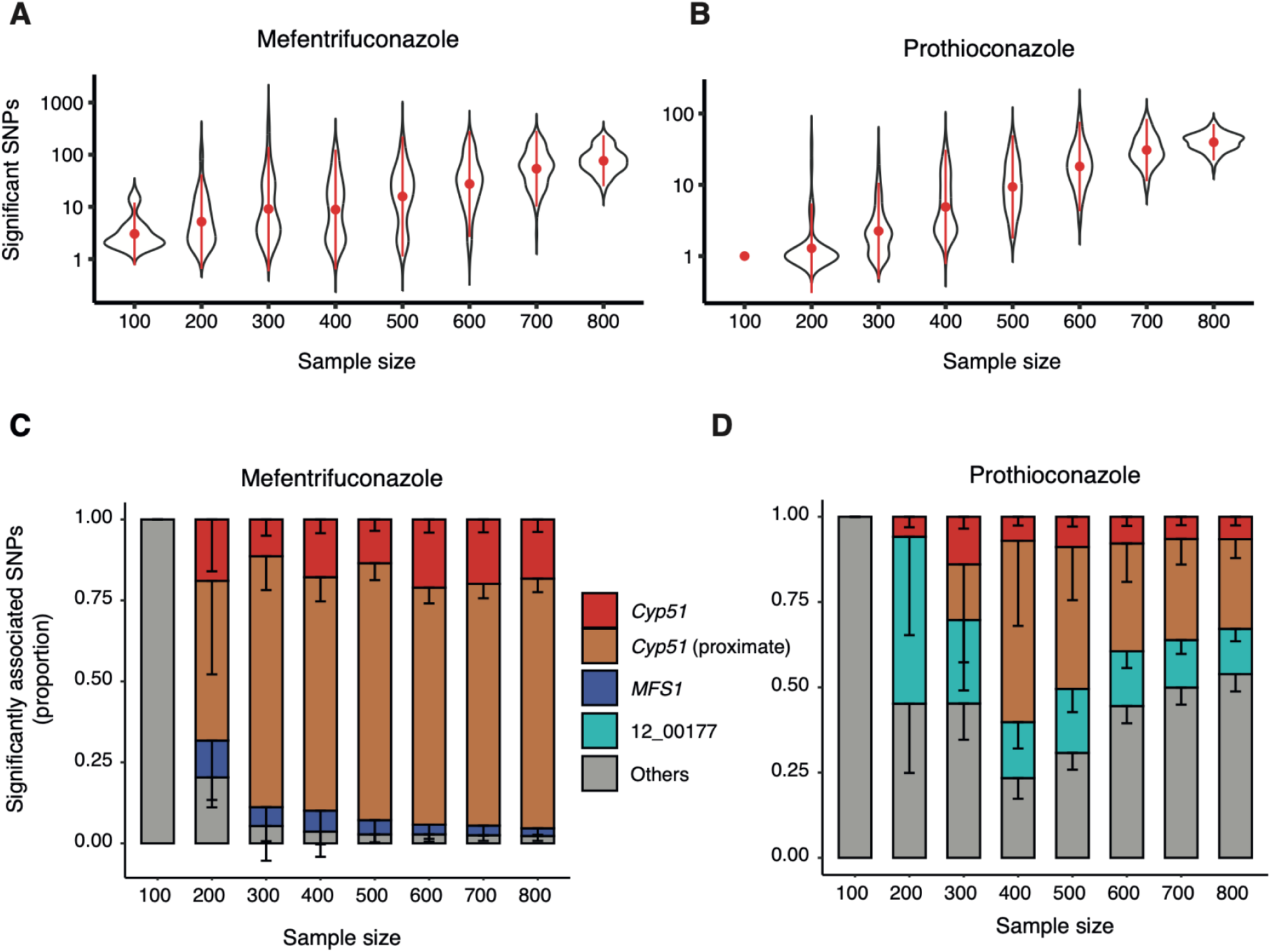
Significant associations based on random subsets of strains assayed for mefentrifluconazole and prothiconazole resistance. (A-B) Average number of highly significant associations for panel subsample sizes (Bonferroni threshold; 100 panel resamplings). (C-D): Significant association counts for associations in proximity to genes with known resistance functions. Color coding indicates the functional class of the closest gene: purple (*Cyp51*), green (others), pink (close to *Cyp51* – from 7_00451 to 7_00459), light blue (MFS1), red (*Cytb*), light green (others), light green (12_00177).

### Optimizing phenotyping assays to improve resistance factor discovery

We first assessed whether fungicide concentration affects the expression of different resistance mechanisms. For this, we analyzed colony growth responses to a range of epoxiconazole concentrations in the medium (10 mg.L^-1^ to 0.01 mg.L^-1^; Figure 4A). Reproducibility of the colony growth assay in the 96-colony spotting format was high comparing replicate plates (*r* = 0.656, *p* < 10^-6^ ; Figure S1; Table S1). Even though single-replicate assays remain unsuitable to assess per-genotype resistance levels with high confidence, a single replicate produces sufficiently robust data for GWAS as shown previously ^5^. At the lowest dose, only 0.2% of all strains were suppressed and GWAS revealed no significant associations. Increased fungicide concentration increased the proportion of susceptible strains (Figure 4B). At concentrations of 0.1 and 1 mg.L^-1^ epoxiconazole in the medium, GWAS identified missense substitutions in the target gene *Cyp51* including Ser524Arg and two SNPs upstream of a gene of unknown function (7_00448; Figure 4 C-D). Starting at 1 mg.L^-1^ fungicide concentration, GWAS identified three synonymous SNPs in the drug exporter gene *MFS1* (Figure 4E). Finally, at the highest dose (10 mg.L^-1^), most strains were repressed and GWAS was unable to map resistance mutations. Across the concentration range, we mapped a total of 266 SNPs significantly associated with epoxiconazole resistance.

**Figure 4:**
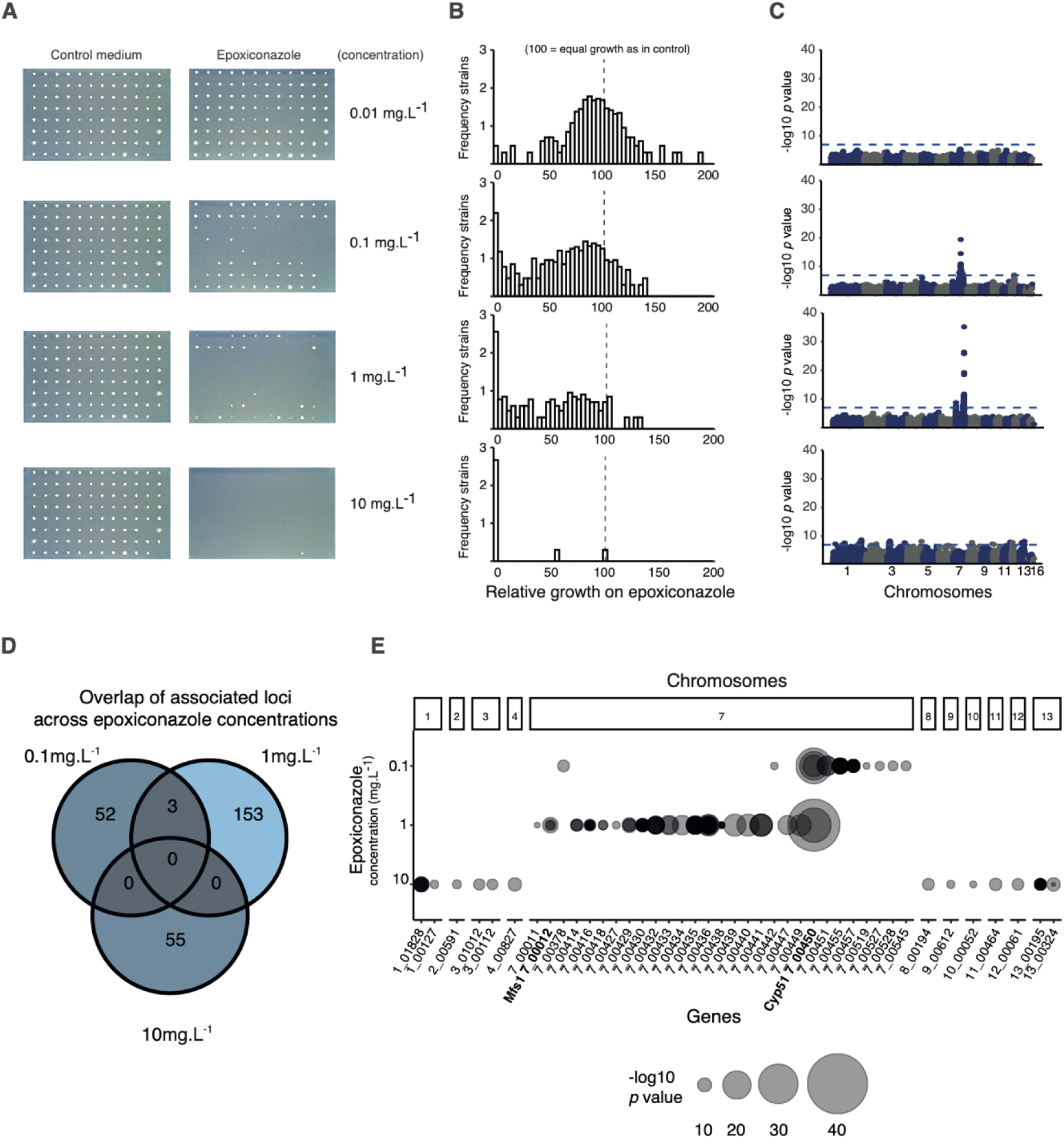
Relative colony growth resistance phenotyping and association mapping. (A): Colony growth in presence of fungicides at 0.01, 0.1, 1, 10 mg.L^-1^. (B): Frequency distribution of relative colony sizes. 0 corresponds to no growth and 100 corresponds to equal growth on control and fungicide medium. C: Manhattan plots showing ignificant associations above the Bonferroni threshold (blue dotted line). D: Venn diagram of SNPs above the Bonferroni threshold. E: Scatter plot *p*-values above the Bonferroni threshold overlapping with a gene detected for resistance assessments in concentrations 0.1, 1, 10 mg.L^-1^ of epoxiconazole and based on relative growth.

Beyond fungicide concentration in the growth medium, colony growth kinetics in response to fungicides can potentially affect GWAS performance. Thus, we assessed colony growth both 7- and 14-days post inoculation (dpi) for four different fungicides. For the QoI azoxystrobin, GWAS retained power regardless of the assay runtime consistently mapping the known mitochondrial *cytochrome b* Gly143Ala resistance mutation (Figure 5A). For the SBI class II fenpropimorph, GWAS identified a single synonymous SNP (660Pro) in the gene 7_00011 upstream of *MFS1* at 7 dpi and three additional SNPs at 14-dpi assay runtime including synonymous variants in *MFS1* (Asn114 and Ile116). No significant associations were observed with the SBI class III fenhexamid regardless of runtime (Figure 5B-C). Finally, we mapped resistance mutations to the succinate dehydrogenase inhibitor (SDHI) carboxin at multiple runtimes. At 7-dpi, we mapped a Glu115Lys missense mutation in a gene encoding an epoxide hydrolase (1_00671) and three mutations in a gene encoding a methyltransferase (8_00208). At the longer runtime, only a downstream mutation near an ABC transporter gene (4_00005) was mapped (Figure 5D). These findings highlight that both variation in fungicide concentration and the colony growth stage have significant impacts on GWAS to map resistance mutations.

**Figure 5:**
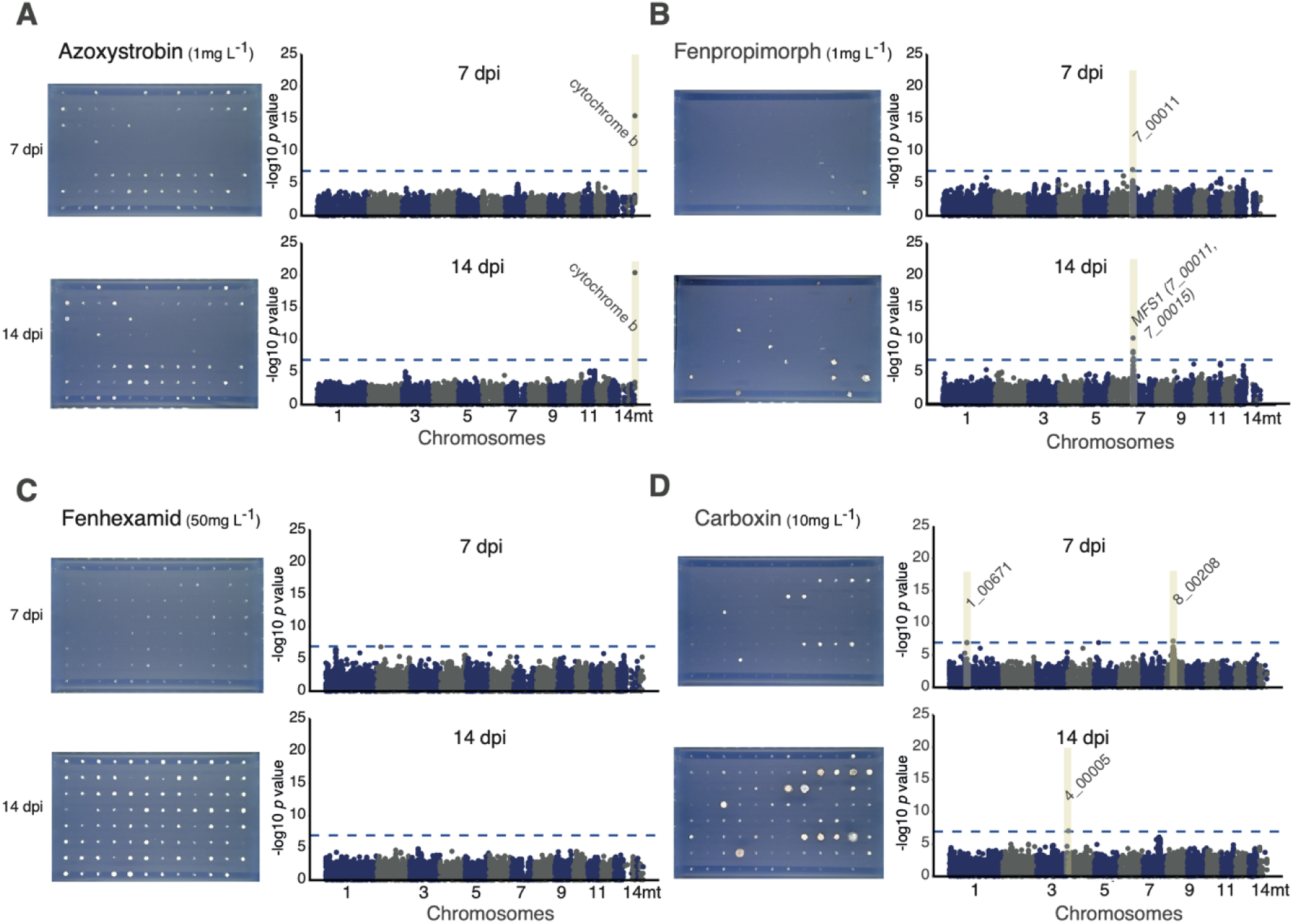
Relative colony growth resistance phenotyping and impact of colony age. (A) Azoxystrobin (1 mg.L^-1^), (B) fenpropimorph (1 mg.L^-1^), (C) fenhexamid (50 mg.L^-1^), (D) carboxin (10 mg.L^-1^). The Manhattan plots show brown rectangles corresponding to regions with SNPs significantly associated with resistance. In the scatterplot, the blue color corresponds to SNPs overlapping with genes significantly associated for measurements at 7 dpi and red colors correspond to 14 dpi data.

### Contributions of complex genetic variants

Structural variants including transposable elements (TEs) and indels impact fungicide resistance of *Z. tritici* ^5,11,25,29^. However, relative contributions of different variant types to overall resistance remains poorly understood. Consequently, we genotyped the European diversity panel using a diverse set of genetic markers beyond SNPs including short indels, copy number variants (CNVs), evidence for newly inserted TEs and a reference-free approach based on k-mers (*i.e.* subsets of short reads). The highly diverse panel segregated a total of 8,536,499 SNPs and 6,606,877 indels. Removing rare variants (<5%), we retained 472,041 biallelic SNPs and 23,959 biallelic indels (Figure S2A-B; Table S3). Next, we assessed GWAS outcomes of each of the different variant classes separately. For this, we included nine fungicide concentrations (0.01–50) across two time points (7 and 14 dpi). Colony growth was assessed in solid media amended with 29 different fungicides and at different timepoints for a total of 43 assay conditions. GWAS produced a total of 901 significant SNPs in a total of 33 assay conditions and identified 186 indels in a total of 28 assay conditions. Furthermore, 66,877 k-mers were significantly associated to variation in fungicide resistance (Table S4). A total of 65.58% of the k-mers could be localized at a unique position in the reference genome. This proportion increased to 98.28% when k-mer locations were searched in the reference-quality pangenome of the species (Figure S2C; Table S5). Recent TE insertion activity creates insertion polymorphism in populations. We identified a total of 2,542 of such TE insertion loci where the inserted TE was present in at least 1% of the panel. Among the loci, 792 TEs were present also in the reference genome and 1,750 TEs were found newly inserted only in strains of the European diversity panel (Figure S2D). Overall, 9% of all TE insertion polymorphisms were significantly associated with fungicide resistance (*n* = 218 loci) dominated by *Copia* and DNA transposons (Table S6). Segmental deletions were assessed in 1-kb windows using a CNV calling pipeline and 68 of such CNVs were significantly associated with fungicide resistance. Such CNVs affected two genes (7_00456 and 5_00703; Table S7). Overall, our assessment shows how genotyping non-canonical variant types such as recent TE activity and reference-free approaches can account for a substantial fraction of total heritable variation in fungicide resistance.

### Establishment of a pan-European fungicide resistance atlas

We performed and integrated GWAS findings for 15 fungicide classes relevant for agriculture. We included compounds with specific recommendation to combat *Z. tritici* in European wheat fields, additional crop protection fungicides, as well as clinical compound classes such as echinocandins (Figure 6A; Table S8). We first performed SNP-based GWAS for 43 distinct assays covering 29 distinct compounds expanding to multiple assay runtimes and compound concentrations. We discovered a total 901 significantly associated SNPs of which 61.1% (*n* = 557) were in coding regions of 146 distinct genes. Of these, 41.8% (*n* = 377) of the SNPs were synonymous variants, 19.1% (*n =* 174) were missense variants and three were altered stop codons. The DMI class included five compounds and accounted for 71.5% of the total SNPs (*n* = 645). Consistent with previous GWAS on DMI resistance using the European diversity panel ^5^, 327 SNPs localized to a 40-kb region near the *Cyp*51 gene encoding the molecular target of azoles. The identification of a large region with highly significant associations is consistent with a selective sweep generating long-range linkage disequilibrium. For three DMIs including cyproconazole, epoxiconazole, and mefentrifluconazole, we identified significant associations for *MFS1* encoding a multidrug exporter channel. Beyond DMIs, *MFS1* variants were associated with resistance to two additional fungicide classes targeting the sterol biosynthesis pathway (*i.e.*, SBI II and SBI IV). Resistance to the four tested SDHIs accounted for 18.4 % (*n* = 166) of the significant SNPs in 18 distinct genes. Isofetamid resistance was mapped to 143 SNPs within ∼10 kb of the *sdhC3* gene encoding the resistance-associated paralog of the SdhC1 subunit ^11^. Resistance to the QoI fungicide azoxystrobin mapped to the well-known Gly143Ala substitution in the mitochondrial *cytochrome b* gene encoding the molecular target ^32^. The assays with the benzylcarbamate-type fungicide pyribencarb revealed resistance mutations in *Cyp51* even though the gene does not encode the molecular target of the compound. Resistance to the QiI fungicide florylpicoxamid was mediated by four genes (1_01890, 3_01116, 9_00415, 11_00307) consistent with non-target site resistance mechanisms. Carbamate and N-phenyl carbamate fungicide resistance were both mediated by a Glu198Ala substitution in the beta-tubulin gene. The Glu198Ala substitution underpinned a strong negative cross-resistance with opposite effects on resistance between the two alleles. Furthermore, the Glu198Ala substitution was significantly associated with resistance to the anilinopyrimidine fungicides pyrimethanil and mepanipyrim. Anilinopyrimidine fungicides were thought, at least in *B. cinerea,* to act on a different molecular target ^33^. Beyond anilinopyrimidine, which is no longer recommended for controlling *Z. tritici*, we also detected resistance mutations for the clinically relevant echinocandin (Table S8).

**Figure 6:**
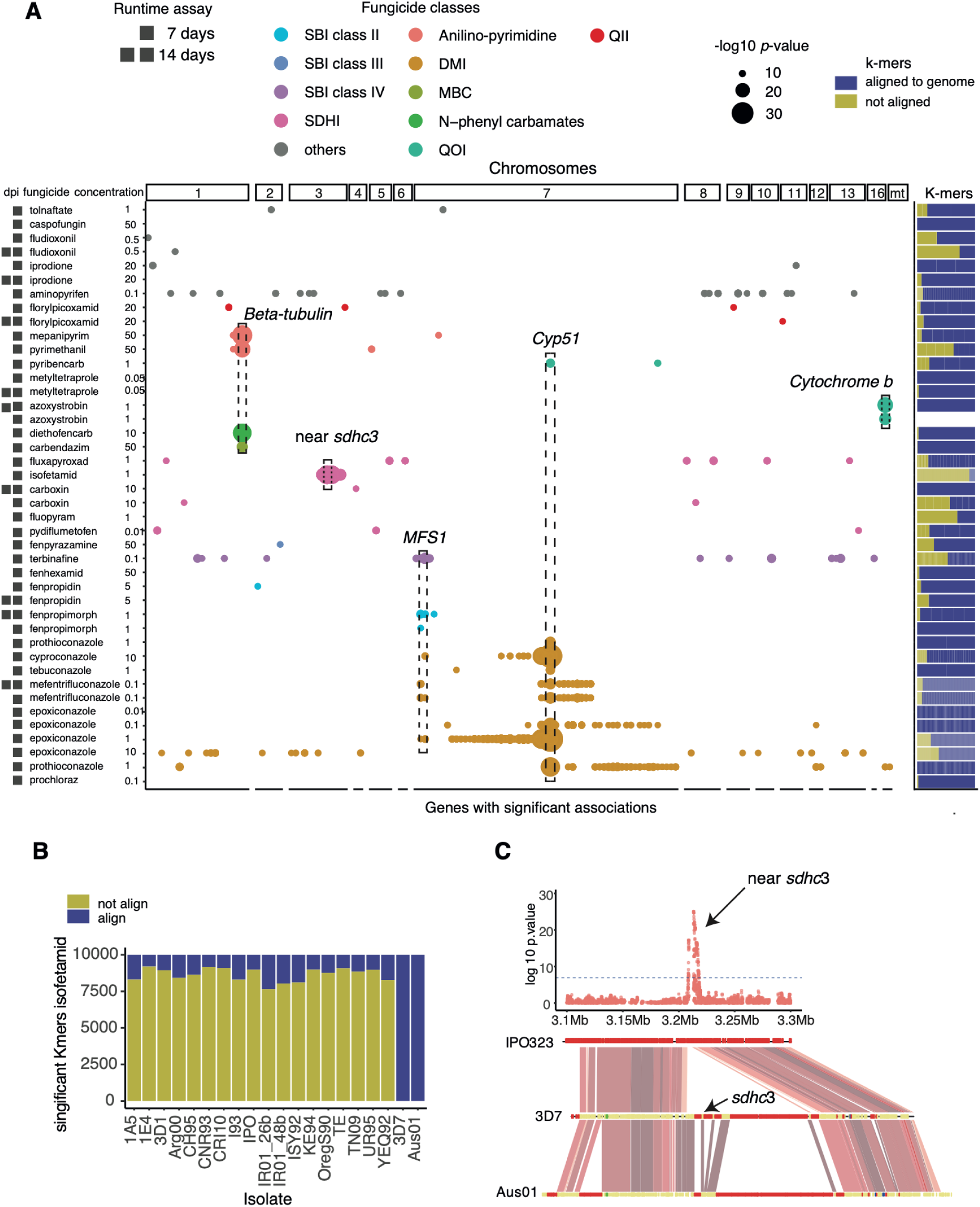
Construction of a SNP and k-mer based fungicide resistance atlas. A: Scatter plot of SNP *p*-values above the Bonferroni threshold overlapping with genes and for the 43 assays based on relative growth. Fungicides are grouped and colored based on classes: SBI class II, SBI class III, SBI class IV, SDHI, anilino-pyrimidines, DMI, MBC, N-phenyl carbamates, QII, QOI, and others (corresponding to PP-fungicides, dicarboximides, carbamothioic acid derivative, echinocandin, 2- aminonicotinate). Proportion of k-mers aligned (yellow) and unaligned (blue) to the IPO323 reference genome. B: Proportion of aligned (yellow) and unaligned (blue) significant k-mers for isofetamid on the species pangenome. In blue, aligning; in yellow, proportion not aligning to the pangenome. C: Manhattan plot for the chromosome 3 of positions 3.1-3.3 Mb and a synteny plot of the same region represented by the reference genomes Aus01 and 3D7 strains highlighting the rearrangement of the locus. In red, TEs; in yellow, annotated gene regions.

### Functional genetics assays to assess GWAS findings

We first analyzed evidence for two resistance mechanisms against fungicide classes not specifically applied to combat *Z. tritici* on wheat. Terbinafine, a class III sterol biosynthesis inhibitor (SBI), is commonly used to treat human fungal infections and was investigated for multidrug resistance in filamentous fungi ^34,35^. However, terbinafine has been used to screen for *Z. tritici* overexpression mutants for the multidrug exporter MFS1 ^26^, which carries a highly plastic promoter sequence ^23^. Both SNP and k-mer based GWAS identified *MFS1* as the top gene associated with terbinafine resistance, with six synonymous SNPs (Figure S3A). Interestingly, a second *MFS* gene (10_00421) was associated with terbinafine resistance through a synonymous SNP (Val531Val) and 86 mapped k-mers (Figure S4 A). To assess contributions of the second *MFS* gene to resistance, we inserted a tetracyclin-repressible promoter upstream of the gene. The promoter is tunable by doxycycline. A promoter swap was also performed at the *MFS1* locus as a control. In addition, we produced a knock-out mutant of the second *MFS* gene to assess whether deletion of the gene would confer oversensitivity to terbinafine (Figure S3B). As expected, overexpression of *MFS1* conferred resistance in media supplemented with terbinafine and epoxiconazole, while doxycycline-induced repression restored sensitivity (Figure S3C, Figure S4B). However, neither overexpression nor the knock-out of the second *MFS* gene altered terbinafine resistance (Figure S5AB). Hence, we found no support to recapitulate the GWAS association in the IPO323 background. Genomic retracing of resistance alleles in the European diversity panel are consistent with the *MFS* genes gaining resistance-associated alleles independently (Figure S6 A).

Next, we focused on structural variants associated with pyrimethanil resistance. K-mer based GWAS identified 396 significant k-mers with 228 aligning exclusively to a reference genome of a North America strain (I93). The unique sequence in the I93 genome to which the k-mers aligned to included an insertion of 1,668 bp on chromosome 1 (Figure S3D). This region encodes the gene *jg.10233* segregating both a functional and a loss-of-function haplotype in the European diversity panel (Figure S7). We hypothesized that the haplotype carrying a functional *jg.10233* confers pyrimethanil resistance. Introducing the *jg.10233* haplotype into the IPO323 genetic background lacking the gene yielded no detectable variation in resistance though. Hence, additional factors may play a role at this locus for pyrimethanil sensitivity, as well as resistance-associated structural variants discovered at other loci (Figure S3E; Table S10).

### Reference-free mapping identifies novel resistance mutation

The reference free k-mer approach expanded the discovery of resistance factors beyond canonical variants mapped to a reference genome. We identified an average of 2699 k-mers associated with SDHI fungicide resistance and an average of 403 k-mers for DMI resistance (Figure 6B). In order to localize the k-mer associations to individual loci in the genome, we performed a k-mer alignment step. For the SDHI isofetamid, an unusually high proportion (89.9 %) of k-mers were not aligning to the reference genome. However, most k-mers could be aligned to a reference-quality pangenome of the species (76- 91% of k-mers aligned; Table S9). The pan-genome based alignment of associated k-mers was identifying a large structural variant encoding an *SdhC3* paralog known to increase SDHI resistance in the field (Figure 6C). Following the successful mapping of the *sdhC3 paralog*, we further investigated resistance mutations inaccessible by SNP-based analyses. The top 20 most significantly associated k-mers for carboxin resistance showed overlaps with the gene encoding *SdhC1*, the primary copy encoding the *SdhC1* subunit of the succinate dehydrogenase complex (Figure 7A; Table S4). Of these k-mers, 16/20 mapped to a Thr79Asn amino acid substitution (Figure 7B), which was missed by the SNP-based GWAS due to a rare variant filtering step (Figure 7A). The Thr79Asn substitution was first detected in 2016 in Germany, followed by a more widespread occurrence across Europe and a notable frequency increase in Ireland ^1^. We functionally validated the impact of the Thr79Asn substitution by generating isogenic mutants for the reference genome strain IPO323. The background was deficient in ku70 (Δku70), which facilitates homologous recombination. Two independent mutants showed significantly increased growth on carboxin-amended media compared to the susceptible IPO323 background. Hence, Thr79Asn is indeed associated with gains of carboxin resistance in Europe and most likely affects the inhibitor binding site of the succinate dehydrogenase complex ^9,11^ (Figure 7D, Figure S8A-C).

**Figure 7:**
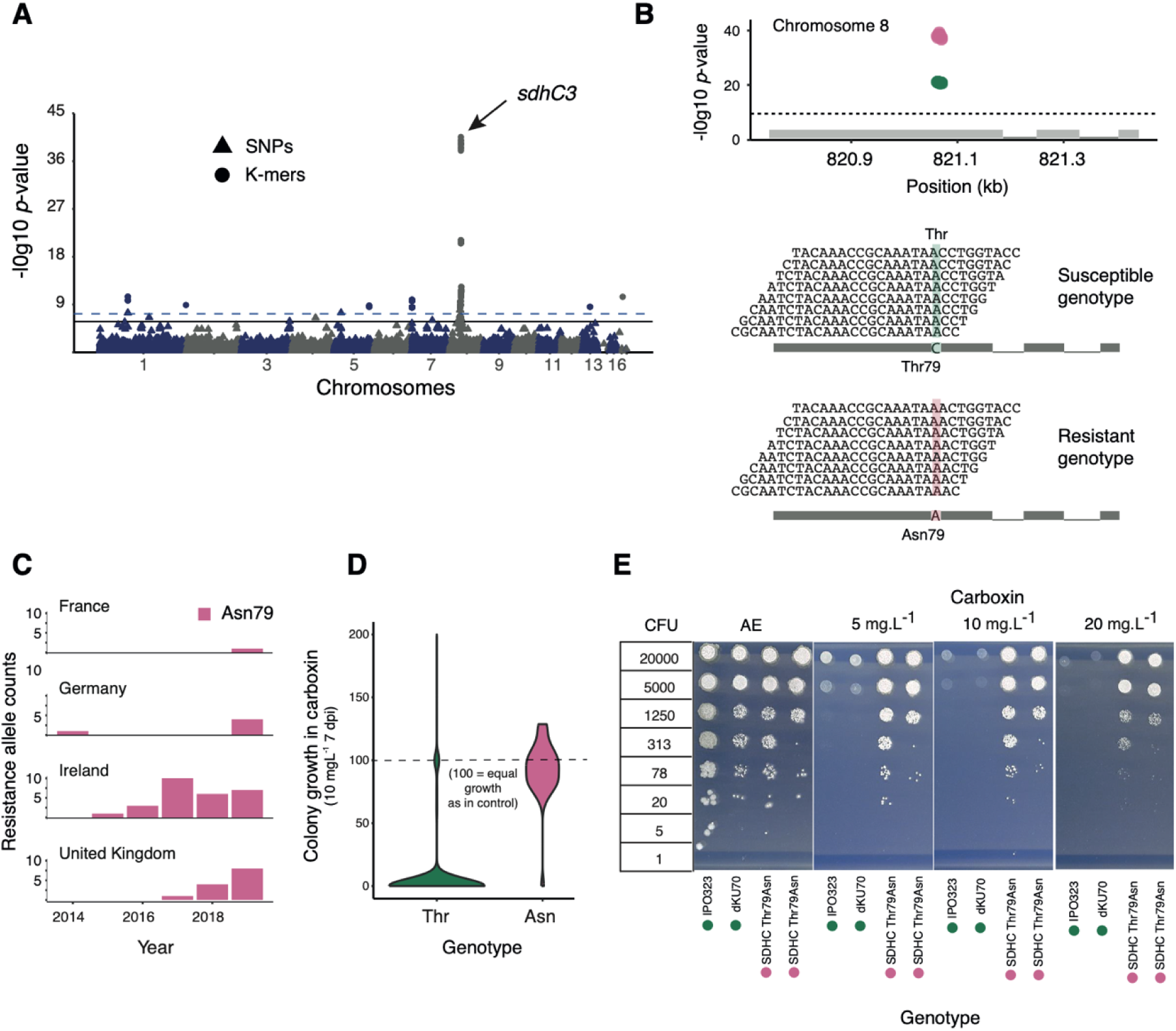
Profiling and functional validation of the Thr79Asn variant in *SdhC1* associated with carboxin resistance. A: Manhattan plot of carboxin (10 mg.L^-1^) resistance associations with k-mers (circles) and SNPs (triangles). B: Focus on the k-mers mapping to the *SdhC*1 gene region at 820.7-821.4 kb on chromosome 8. Multiple sequence alignment of k-mers matching either the Thr residue (in green) or Asn (in red) at the codon position 79. C: Frequency of the Asn residue across the European diversity panel. D: Violin plot comparing relative growth values in strains with Thr and Asn. E: Functional validation of mutants shown on a spotting assay including IPO323 (wild type), dKU70 (green dots) and SDHC_Thr79Asn (red dots) on AE media amended with carboxin at 5, 10, 20 mg.L^-1^.

### Negative cross-resistance in beta-tubulin

The fungicide resistance atlas revealed a negative cross-resistance associated with the Glu198Ala substitution in the beta-tubulin gene. Encoding Glu at position 198 is associated with higher resistance against diethofencarb (a methyl-benzimidazole carbamate class member). Encoding Ala at the same position confers resistance to carbendazim (N-phenyl carbamate class) (Figure 9C) ^36^. The negative cross-resistance was previously characterized by showing that the Glu198Ala substitution confers resistance to carbendazim and sensitivity to diethofencarb ^37^. We found that Glu198Ala also mediates resistance to the anilinopyrimidine fungicide class (such as pyrimethanil and mepanipyrim fungicides). The anilinopyrimidine fungicides have not been used for *Z. tritici* control or for the control of wheat pathogens in general. We observed a strong antagonistic growth response for strains growing either well on carbendazim (10 mg.L^-1^) or diethofencarb (10 mg.L^-1^) (Figure 8A), as well as between carbendazim (10 mg.L^-1^) and pyrimethanil (50 mg.L^-1^). Strains more susceptible to pyrimethanil tended also to be more susceptible to diethofencarb (Figure 8B-C), however the correlation in growth responses was not significant (*p* = 0.49; Figure 8C). We used molecular genetics to assess the impact of the point mutation associated with pyrimethanil resistance. For this, we generated isogenic lines of the IPO323 Δku70 background swapping the beta-tubulin Glu198 for Ala (GAG to GCG) (Figure 8D). The resulting mutant with the Ala substitution exhibited resistance to carbendazim, and susceptibility to both diethofencarb and pyrimethanil. Hence, the negative cross-resistance predicted by the fungicide resistance atlas indeed replicates in functional assays.

**Figure 8:**
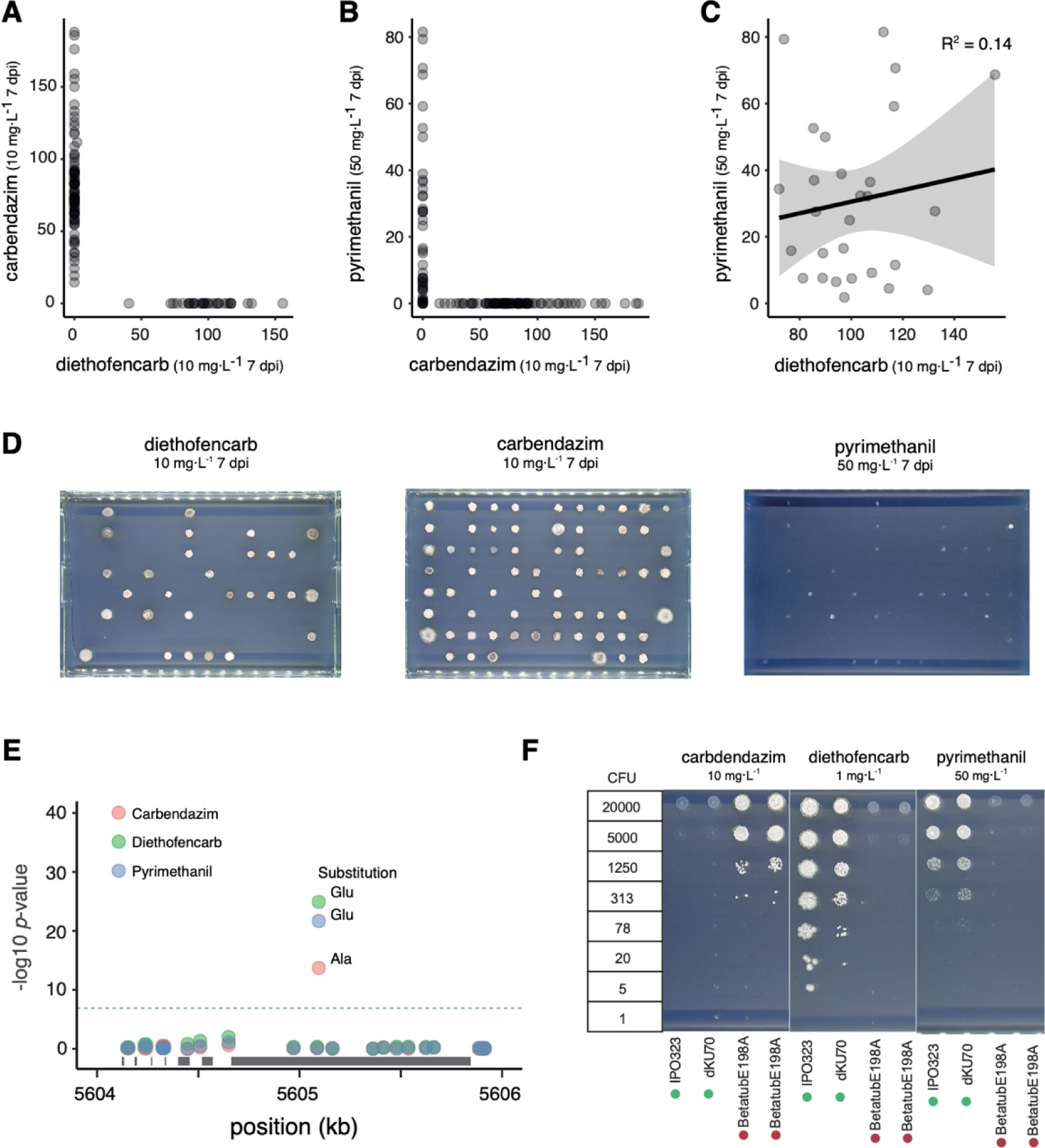
Profiling and functional validation of the Glu198Ala variant in *beta-tubulin* associated with pyrimethanil, diethofencarb, and carbendazim resistance. A: Correlation of relative growth values between carbendazim and diethofencarb. B: Correlation of relative growth values between pyrimethanil and carbendazim. C: Correlation of relative growth values between pyrimethanil and diethofencarb. D: Fungal colonies in the presence of pyrimethanil (50 mg.L^-1^), diethofencarb (10 mg.L^-1^), and carbendazim (10 mg.L^-1^). E: Zoomed Manhattan plot of significant SNPs for pyrimethanil, diethofencarb, and carbendazim, highlighting the *beta-tubulin* gene region (506.4–506.6 kb) on chromosome 1. Each significant SNP is labeled with the resistance-associated allele (Thr or Asn). F: Functional validation of the mutants shown on the spotting assay with IPO323 (wild type), dKU70 (green dots), and SDHC_Thr79Asn (red dots) on AE medium amended with pyrimethanil (50 mg.L^-1^), diethofencarb (1 mg.L^-1^), and carbendazim (10 mg.L^-1^).

## Discussion

A comprehensive understanding of genetic factors contributing to emerging fungicide resistance is crucial for effectively managing fungicide deployment and development of new compounds. This study introduces a high-throughput pipeline capturing continent-wide genetic diversity of a major crop pathogen. The genomic survey revealed a complex landscape of resistance against dozens of compounds. Complex genetic variation makes important contributions to overall resistance. We refined association mapping studies with analyses of structural variation and confirmed some key predicted resistance mutations using functional assays.

Diversity panel size, genetic and phenotypic variability affect the power of GWAS analyses ^19^ ^20,38^. The evaluated European diversity panel comprises 1420 strains sampled in a hierarchical manner ^5^. We show that even a small fraction of the panel could represent a large proportion of the continent-wide pathogen diversity. However, small panel sizes would have lacked sufficient mapping power to detect most resistance loci discovered in the largest panel sizes. Earlier population genetic analyses and monitoring programs have highlighted the heterogeneous distribution of resistance across Europe ^15,39^. We found direct evidence for this heterogeneity by contrasting panel subsets split along either an east-west axis or segmented by collection years. We also found that the discovery threshold to robustly discover variants was highly dependent on the identity of the fungicide. This is most likely explained the magnitude of individual mutational effects on resistance but also by mutation frequencies in the panel. The mapping power analyses also revealed that prothioconazole resistance mapped to the mitochondrial *cytochrome b*, beyond the expected molecular target gene or the multidrug resistance factors. A possible explanation for the association may be high geographic congruence of DMI and QoI exposure and highly correlated resistance gains. Fungicide combination treatments in agriculture may lead to co-selection for dual resistance and, hence, genetic correlations. Furthermore, the multigenic basis of prothioconazole resistance may be related to the conversion step of prothioconazole into prothioconazole-desthio by oxidation in fungal cells, essential for Cyp51 inhibition and antifungal activity ^40^. Finally, loci associated with strong resistance (*i.e. Cyp51* substitutions) were detected already at low fungicide doses. Non-target mechanism such as MFS1*-*mediated multidrug resistance, required exposure to higher fungicide doses in the assays. This is consistent with a model of non-target resistance mechanisms making only a perceptible contribution to resistance in backgrounds encoding also target resistance mechanisms. We expect that at low fungicide dose, target resistance mutations overwhelm variation in growth responses. However, at high dose, only genotypes with target resistance are able to grow and that this baseline resistance is enhanced by non-target resistance and becomes accessible by association mapping. The substantial improvements of fungicide resistance GWAS through assay optimization highlights the large potential of pathogen diversity panels to uncover resistance mechanisms.

We show that genotyping cryptic genetic variation is crucial for the mapping of fungicide resistance loci in crop pathogens. The established resistance atlas comprises a wide range of polymorphisms including SNPs, k-mers positioned in the genome, as well as indels, TE insertion variants and CNV loci. The atlas confirmed previously discovered substitutions in *Cyp51* including Ser524Thr, Ile381Val, Ser188Asn,Val136Phe and Asp134Gly ^41,42^, Gly143Ala in *cytochrome b* ^13^, Glu198Ala in beta-tubulin, as well as genetic variation associated with the multidrug resistance factor MFS1 ^25^. Non-canonical genetic variants beyond SNPs were known to contribute to pathogen trait variation including melanin production, asexual reproduction and fungicide resistance ^26,43,44^. Here, we significantly expanded the repertoire of loci contributing to such trait variation. In particular, recent TE activity is having a considerable impact on fungicide resistance with a tenth of all high-frequency (>1%) polymorphic TE insertion loci showing an association. A hotspot for resistance gains mediated by TEs is the promoter region of *MFS1* ^25,26,45^. We identified previously unknown contributions by TEs such as fluxapyroxad resistance as well as confirming the impact of known TE insertions ^26^. Our work shows that expanding genotyping approaches to cover cryptic variants beyond single reference genomes was critical. While approximately two-thirds (∼65%) of the associated k-mers aligned to the reference genome, expanding the mapping to a reference-quality pangenome resolved nearly all k-mer localizations ^29^. Hence, reference-free GWAS is an essential and powerful tool for identifying standing resistance in the pathogen.

Connecting variant associations to molecular mechanisms is challenging but provides essential information on the biological significance of emerging resistance. We prioritized a set of loci identified through distinct GWAS approaches. We confirmed the Thr79Asn substitution mapped by k-mer association in the *SdhC1* gene ^46^, which likely interferes with inhibitor binding. Furthermore, we recapitulated the predicted negative cross-resistance effects of a single substitution in the beta-tubulin gene. The mutation confers resistance to the anilinopyrimidines pyrimethanil, which may be a species-specific property. In *B. cinerea*, the activity of anilinopyrimidines fungicides is reversed by the addition of methionine and field resistance is driven by mutations in at least two genes involved in mitochondrial processes. Therefore, anilinopyrimidines in *B. cinerea* are thought to primarily target a mitochondrial function connected to methionine biosynthesis ^33^, whereas high doses affect tubulin function in *Z. tritici*. We failed to replicate the GWAS discovery of 10_00421, an MFS gene associated with terbinafine resistance. Possible explanations include unaccounted for epistasis, where the effect of the resistance mutation might depend on additional mutations not accounted for in the genetic background. Similarly, how an associated structural variant confers pyrimethanil resistance could also not be retraced. Designing molecular genetic assays replicating GWAS findings are notoriously challenging as associations may be contingent on the genomic background. However, more efficient assay designs and the systematic screening of multiple genomic backgrounds for hypothesis testing should reduce this challenge.

Overall, our work establishes a fungicide resistance atlas recapitulating the continent-wide emergence of resistance to a comprehensive set of fungicide classes, including the most frequently applied DMIs, SDHIs and QoIs. We provide concrete guidance how pathogen diversity panels should be constructed to unravel resistance emergence in the field. In particular, we show how to mitigate mapping power limitations, and to what extent geographic and temporal sampling is relevant. Systematic reference-free genotyping of complex variants is crucial to capture the full spectrum of resistance gains. Building a pangenome-informed atlas of emerging resistance benefits both the discovery of resistance mechanisms and guides towards more sustainable applications of compound mixtures.

## Material and methods

### European diversity panel

For resistance monitoring purposes, Syngenta Inc. sampled pathogens from commercial and trial sites across Europe. The fungicide resistance monitoring efforts for *Z. tritici* led to a collection of 8607 strains. Strains were selected from the collection based on a hierarchical clustering approach grouping sampling locations into 100-km radius areas as described in detail previously (REF Chap 1). Within each area, the collection was balanced over the 2005-2019 sampling years based on availability. The final European diversity panel included 1420 strains from 27 different countries ^5,21^ (Table S1). Estimation of fungicides application were based on the yearly sale in kg across the continent. The information was obtained based on the average fungicide application within the time range 2011-2021^30^. Wheat production per country was obtained by FAO and was average in tons between 2017-2019. Maps were plotted in ggplot using the package (‘rnaturalearth’) ^47,48^.

### Purification of the European diversity panel

All strains included in the European diversity panel were collected from infected leaves across Europe, preferably with visible pycnidia, wrapped in dry paper towels, and packed into a paper envelope for shipment. Upon arrival in the laboratory, samples were labeled with a unique code, and the arrival date was recorded. The leaves were dried, wrapped in fresh paper towels if necessary, and stored at 4°C until further processing. Leaves with symptoms were cut into 2 cm pieces, surface-sterilized in 2% bleach for 2 minutes, and rinsed with sterile distilled water. Leaf cuts were placed on wet filter paper in Petri dishes (1.3 mL water for 9 cm dishes) and incubated at 20°C for 24 hours. Single strains were picked from cirri under a binocular microscope and transferred to V8 agar plates with antibiotics. Plates were incubated for 4-7 days at 20°C. To ensure that only a single genotype was collected per sample, a single spore isolation step was performed for every sample. Single colonies were subcultured on fresh V8 plates and incubated under the same conditions for an additional 7 days. Each isolate was stored independently in a cryovial preserved in liquid nitrogen. Fresh cells harvested from V8 plates incubated at 18°C for a period of five days were used as inoculum for all experiments. Media recipes used in this study have been summarized (see Table S11).

### Solid culture assays for fungicide sensitivity determination

We assessed fungicide sensitivity using relative colony size estimates on solid medium (fixed fungicide concentration). The European diversity panel strains stored at −80C° were arrayed in 96-well cryo-stock plates in skim milk with 20% glycerol (used as mother array plates). From the mother array plates, the strains to be tested were transferred using a 96-floating pin replicator tool (408FS2AS, V&P Scientific Inc.) to 96-well flat bottom plates (model 3370, Corning Inc.) pre-filled with 100μl YPD liquid medium and left to grow at 18°C for 7 days to reach a growth plateau. Spore suspensions were not further standardized and directly spotted using a Rotor HDA (Rotor+; Singer Inc., Watchet, UK) with sterile pins (RePads) (Singer Inc., Watchet, UK) onto AE agar media plates (PlusPlates) amended or not with fungicides. The fungicides were dissolved in DMSO and then mixed with AE-agar to achieve a final concentration of 1% DMSO. Five different fungicide concentrations were applied to achieve final concentrations of 50, 10, 1, 0.1, and 0.01 mg.L^-1^ per plate. Single concentrations were then selected for further GWAS studies. Control plates only contained AE-agar with 1% DMSO. Spotted AE-agar plates were incubated at 20°C for 7 days in the dark before imaging. Image capture was performed using a Phenobooth (Singer Inc.), and image analysis performed with the Phenosuite (version 2.21) software package (Singer Inc.) by comparing paired growth areas of colonies grown on control plates without fungicide (DMSO controls) with fungicide-amended plates. The relative growth area on amended media (*area*_e_) relative to controls (*area*_c_) was estimated using the following formula:

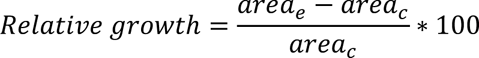

Plates with signs of contamination were excluded. Correlation between two control replicates was estimated using the *area*_c_ at 7 dpi in AE media in absence of fungicides (Table S12). Plates with >2x more growth on fungicide-amended plates *vs.* controls were excluded as well. Relative growth estimates and assayed fungicides are reported in Table S1 and Table S2. Fungicide sensitivity of the transformants was determined on solid medium (fixed fungicide concentration). For the spotting assay the suspended spores were spotted on AE media amended with 1% of a 100× DMSO solution of the active ingredients to pre-melted AE-agar. AE media was supplemented with 20,5,2.5 mg.L^-1^ of terbinafine and epoxiconazole and each experiment was performed with and without doxycycline (30 mg.L^-1^). We supplemented AE media as well with 10 and 50 mg.L^-1^ of carbendazim, 1 and 10 mg.L^-1^ of diethofencarb, 20, 50 mg.L^-1^ of pyrimethanil and 5,10 and 20 mg.L^-1^ of carboxin. Pre-culture of the inoculum and fungicide sensitivity tests were performed as following previously described procedures^9^.

### DNA extraction, Illumina sequencing, and SNP calling

Whole-genome sequencing data was previously described ^5,21^. For 1134 strains high-quality genomic DNA was extracted using the DNeasy Plant Mini Kits (Qiagen Inc.) following the manufacturer’s instructions. Paired-end sequencing of 250 cycles with an ∼500 bp insert size was performed by Novogene Inc. using the Illumina NovaSeq 6000 platform. We used Trimmomatic v.0.39 to trim low-quality sequencing reads and remove adapter contamination in each isolate ^49^. Filtered sequences were aligned to the *Z. tritici* reference genome IPO323 ^50^ using Bowtie2 v.2.3.3 ^51^. The Genome Analysis Toolkit (GATK) v.4.0.1.2 ^52^ was used for single nucleotide polymorphism (SNP) calling and variant filtration. The GATK HaplotypeCaller was run with the command -emitRefConfidence GVCF and - sample_ploidy 1. Joint variant calling was performed using the tool GenotypeGVCFs merging HaplotypeCaller gvcfs produced for an additional 283 from a previous study ^21^ with the option -maxAlt 2 ^53^. To investigate the impact of sample size on the number of detected alleles, we randomly subsampled the European collection 100 times. The number of strains included in each subsample varied from 100 to 1000. Vcftools v0.1.15 ^54^ was used to perform the random subsetting. To estimate the total number of SNPs based on the annotation of the *Z. tritici* 2015 genome ^55^, we used SnpEFF ^56^ with the “-csvStats”. Finally, the total number of synonymous and missense point mutations was normalized based on the corresponding gene length (see Table S13).

### SNP-based genome-wide association mapping and bootstrapping

For SNP-based GWAS on the Europe diversity panel, we retained SNPs with a genotyping call rate of ≥90% and a MAF ≥5% resulting in a final set of 472,103 biallelic SNPs (Table S1). Traits for GWAS included the relative growth estimates for 43 traits (Table S2). We tested 15 fungicide classes with an over representation of DMI, SDHI, QOI, QII, classes applied in *Z. tritici.* We accounted for relatedness by constructing a genetic relatedness matrix (GRM) among strains using all genome-wide SNPs with the option “-gk 2” in Gemma ^57^. Thus, all the associations were performed using a univariate linear mixed model (MLM+K) where K is the GRM as a random effect:

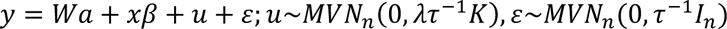

*y* represents a vector of phenotypic values for *n* individuals; *W* is a matrix of covariates (fixed effects with a column vector of 1), *α* is a vector of the corresponding coefficients, including the intercept; *x* is a vector of the genotypes of the SNP marker, *β* is the effect size of the marker; *u* is a vector of random individual effects; *ε* is a vector of random error; *τ*-1 is the variance of the residual errors; *λ* is the ratio between the two variance components; *K* is the *n* × *n* GRM and *I_n_* is an *n* × *n* identity matrix and *MVN_n_* represents the multivariate normal distribution. We applied a stringent Bonferroni threshold (*α* = 0.05; *p* = *α* / total number of SNPs) to identify most robust SNP associations. Significant SNPs were annotated using snpEff v5.0e ^56^. We provide links to all association mapping outcomes in Table S14. To investigate the impact of geographic and temporal distribution on the number of detected resistant alleles, the European diversity panel was split in strains collected from the West and East of Europe after 2016. For the temporal distribution, we split the strains before and after 2016. T-test between relative growth values of West and East and before-after 2016 strains were performed using the command t.test built-in function in R. Subsequently, the results were visualized through figures generated in R ggplot2 and adjusted in Illustrator ^48^. In the Manhattan plot for *p*-value within 0.1 and 0.01 we retained only 1 out of every 30 rows, for *p*-value within than 0.2 and 0.1 we retained only 1 out of every 300 rows and for *p*-value greater than 0.2 we retained only 1 out of every 400 rows for visualization purposes. To investigate the impact of sample size on the number of detected resistant alleles, the number of strains included in each subsample varied from 100 to 800 strains. Vcftools v0.1.15 ^54^ and gemma were used to perform the random subsetting and the GWAS. SNPEFF ^56^ was utilized to annotate the SNPs and genes based on the *Z. tritici* 2015 genome annotation ^55^ (See Table S15 and Table S16).

### K-mer based genome-wide association mapping

We performed k-mer based GWAS on the same 43 traits used for SNP-based GWAS following a previously described approach ^58^. We used k-mers of 25-bp length as suggested for small genomes such as *Z. tritici* following previous studies ^29^. Quality-filtered sequencing reads (see above) of 1406 strains from the European diversity panel were used for k-mer screening. K-mers were counted using a two-step process: The canonization involved treating each k-mer and its reverse complement as equivalent. In contrast, non-canonization treats the k-mer and its reverse complement as distinct entities. A k-mer and the reverse complement are supposed to have the same chance to appear in fastq files. However, bias can favor the presence of one of the two forms for some k-mer. Therefore, k-mers were then filtered based on two criteria: (1) a k-mer must appear in both its canonized and non-canonized forms in at least 5% of the strains, and (2) it must appear in both forms in at least 20% of the strains in which it was found. A GRM was estimated with EMMA (Efficient Mixed-Model Association) as an identity-by-state (IBS) matrix under the assumption that each k-mer has a small, random effect on the phenotype. GWAS was performed by using an LMM+K model in Gemma with a likelihood ratio test to determine p-values. Beta estimation and significance testing were performed using the patched version of kmers_gwas.py, incorporating the recommendations outlined (https://github.com/voichek/kmersGWAS/issues/53 and https://github.com/voichek/kmersGWAS/issues/91). A k-mer was significant when the p-value passed the permutation-based threshold as described by Voichek and Weigel (2020). For all k-mer association mapping see Table S17. We attempted to map all significant k-mers for each trait to the reference genome using the short-read aligner bowtie v1.2.2 ^51^ with the command “-a --best --strata” and retained the k-mers with a unique alignment (see Table S4). We used the center position of the mapped k-mer to the reference genome as a coordinate to inspect nearby features using BEDtools v2.31.0 ^59^. Finally, the proportion of significant k-mers aligning and not aligning across the 19 pangenomes ^22^, per each fungicide, was obtained with bowtie v1.2.2, with the command “-a –best –strata” (Langmead and Salzberg, 568 2012) (see Table S9). The proportion of significant k-mers aligning and not aligning to I93, UA005, I93 jg.10233 and SV_UA005.jg.10233 regions were obtained with bowtie v1.2.2, with the command “-a –best –strata”. We provided each k-mer association mapping file (see Table S14).

### TEs detection and association study

We performed TE based GWAS on the same 43 traits used for SNP-based GWAS. TE annotation was performed with ngs-te-mapper2 ^60^. This is a three-stage pipeline. In the first stage the sequencing reads are queried against a library of TE sequences, for which we used the TE consensus sequences obtained from 20 fully assembled genomes of 19 global strains ^22^. The “junction reads” that align both on a TE consensus sequence and on the flanking, genome are used to determine the site of insertion of reference and non-reference TEs. Non-reference TEs were categorized as reference TEs when located within proximity (up to 100 bp) to a reference TE of the same family. This adjustment aimed to mitigate potential inaccuracies in detection while enhancing the overall frequency of detected TEs. Consequently, insertions of the identical element within a narrow window were treated as a singular insertion for subsequent analyses. For the association analysis, we set the maf to 1% and we used the relatedness matrix (GRM) estimated from european vcf the genome-wide SNPs using the option “-gk 2” in GEMMA ^57^. A summary of the significant TEs is in Table S6. We provided each TE association mapping file (see Table S14).

### CNV genotyping

We performed CNV-based GWAS on the same 43 traits used for SNP-based GWAS. We trimmed the Illumina raw reads with Trimmomatic v.0.32 ^49^ and mapped to the *Z. tritici* (IPO323) reference genome with Bowtie2 v.2.4.0 very-sensitive-local option. To define copy number variation (CNV) to the dataset (n=1420 samples) we used GATK germline copy number caller v.4.1.9.0^52^ in case mode against a previously constructed GATK CNV cohort model built from 1109 *Z. tritici* genomes^61^. CNV interval was set to 1000 bp windows with no overlap. We filtered for GC content in windows (min=0.1 and max=0.9), as well as extremely low and high read counts (--low-count-filter-count-threshold = 5, -- extreme-count-filter-minimum-percentile = 1, --extreme-count-filter-maximum-percentile=99). We used a prior table for chromosomal ploidy to assign prior probabilities for each ploidystate. Finally, we called CNV genotypes using the Determine Germline Contig Ploidy, Germline CNV Caller and Post process Germline CNV Calls functions in case mode. CNV GWAS analysis involved preprocessing the data by removing duplicated chromosomes, converting CNV events into four categories (more than one copy, one copy, no copy, or missing), and identifying and removing the least frequent allele at each position, retaining only the most frequent allele. The association analysis was conducted using GEMMA ^57^ with a minor allele frequency threshold of 5% and with GRM constructed from genome-wide SNPs using the “-gk 2” option. A summary of the significant CNVs is in Table S7. We provided each CNV association mapping file (see Table S14).

### Indels-based genome-wide association mapping

For Indels-based GWAS on the Europe diversity panel, we retained Indels with a genotyping call rate of ≥90% and a MAF ≥5% resulting in a final set of 23,959 indels. Traits for GWAS included the relative growth estimates for 43 traits (Table S2). We tested 15 fungicide classes with an over-representation of DMI, SDHI, QOI, QII, classes applied in *Z. tritici.* We accounted for relatedness by constructing a genetic relatedness matrix (GRM) among strains using all genome-wide SNPs with the option “-gk 2” in GEMMA ^57^. Thus, all the associations were performed using a univariate linear mixed model (MLM+K), where K is the GRM as a random effect. A summary of the significant Indels is in (Table S18). We provided each indel association mapping file (see Table S14).

### Mutant selection criteria

Locus refinement was conducted employing stringent selection criteria encompassing *p*-value significance, effect size, gene functionality, and mutation type (intragenic vs. intergenic). The significance threshold for the *p*-value was determined by adhering to Bonferroni’s correction. Additionally, we focused on selecting mutations with a beta (>=2), indicative of a twofold effect. Missense point mutations were accorded higher priority over synonymous mutations. Moreover, emphasis was placed on previously characterized functions, such as MFS, and SDH subunits, while structural variants were given preferential consideration. The list of the mutants is in Table S19.

### Molecular methods for transformations

All the genes were synthesized by GENEWIZ (Suzhou, China) and were inserted in the multisite pNOV2114 Hyg_gateway. GenBank files are available in Supplementary File S1. To generate the KO of 10_00421 (KO_ Zt09_10_00421), we designed the final plasmid Hygr_pNOV2114_KO_G10877_MFS. The pNOV2114 backbone was supplemented with a region 799bps upstream the 5’ of 10_00421 gene and 2274 downstream 3’ of the 10_00421 gene. We substitute the 10_00421 gene with a short terminator, followed by the constitutive promoter TrpC ^62^, and with the hygromycin resistance cassette (see Figure S3B for a graphical overview and supplemental file Table S19 with plasmids sequences). For the structural variants SV_I93.jg.10233 and SV_UA005.jg.10233 we inserted in the IPO323 background with a region of 1560 bps at position 5571672 on chromosome 1. The flanking region included 977 bps upstream and 1000 bps downstream, the position 5571672 on chromosome 1. The Hygr_pNOV2114_jg10233_I93 and Hygr_pNOV2114_jg10233_UA005 included a short terminator, followed by the constitutive promoter TrpC ^62^, followed by the hygromycin resistance cassette and, respectively the structural variants of I93 (1556 bps) and of UA005 (1560 bps). For generating the expression constructs under the control of a tetracycline-repressing promoter we designed a fragment containing the full Tet repressor expression cassette followed by operator sequences fused to mfaI minimal promoter upstream the starting codon of *Mfs1* (Hygr_pNOV2114_OE_G07914.1_MFS1) and 09_00421 (Hygr_pNOV2114_OE_G10877_newMfs). For the Single swap (SW) point mutation we selected a region of 1888 bps encompassing the *beta-tubulin* (pNOV2114_SW_G02218_BetaTUB) gene and a region of 2017 bps encompassing the *SdhC1* gene (pNOV2114_SW_G09103_SDHC). All entry and subsequent binary plasmids were transfer to *A. tumefaciens*. *Z. tritici* transformation was performed as described previously ^5,63^. *Z. tritici* transformants were selected on Hygromycine, Carbendazim (AE + 10 mg.L^-1^) or Carboxin (AE + 10 mg.L^-1^) depending by the selection cassette. All clones were then validated by PCR and three combination of primers were tested (Table S20). For the single-point mutation, a Sanger sequencing step was included to confirm the cloned fragment.

### AlphaFold2

We further investigated the 3D structure of SDHC1 using AlphaFold2 ^64^. We imputed the sequences SDHC1 (08_00228) to predict the 3D structure. Local Distance Difference Test (lDDT) scores were employed to assess the confidence in amino acid positioning within their local structural environments. While the first few amino acids generally exhibit lower lDDT scores, the score at position 79 was high (approximately 100), indicating a high degree of confidence in its predicted location. However, it’s important to acknowledge that the lDDT score at this position could be influenced by the low coverage of SDHC1 sequences, particularly at the beginning of the protein, within the AlphaFold2 database.

## Acknowledgements

We thank Nadja Lindenberger, Regula Frey, Salvatore Accardo and the Syngenta monitoring team for collecting the strains analyzed in this work. Ursula Oggenfuss provided feedback on the manuscript.

## Declarations

### Author contributions

GP, GS and DC conceived the study, GP performed the research and analyzed the data. GP, DF, DE and CD performed analyses. SFFT provided materials. TB and SMT provided datasets. GS and DC supervised the work and acquired funding. GP, GS and DC wrote the manuscript.

### Data availability

Genome sequencing data is available on the NCBI Sequence Read Archive (accession numbers are provided in Supplementary Table S1). Additional datasets are made available in supplementary materials.

