## Supplementary Figures for "High-throughput discovery of emerging antifungal resistance in crop pathogens"

Puccetti *et al.*

A

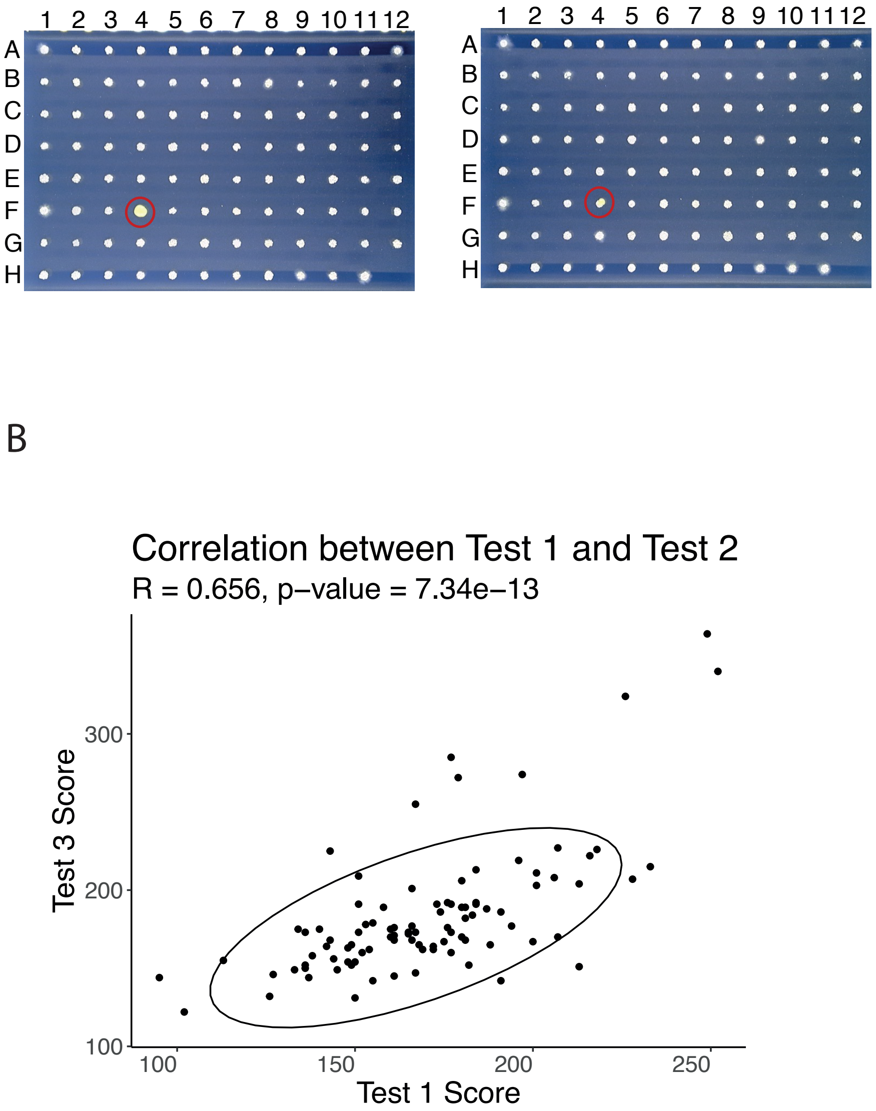


**Figure S1**: Correlation of colony growth across two independent replicates for 95 strains. (A) Representative images of replicate 1 and replicate 2 from an assay performed under control conditions in the AE medium. The colony circled in red was excluded. (B) Correlation of colony area measurements across the two replicates and Pearson's correlation statistics. The full dataset is reported as Table S12.

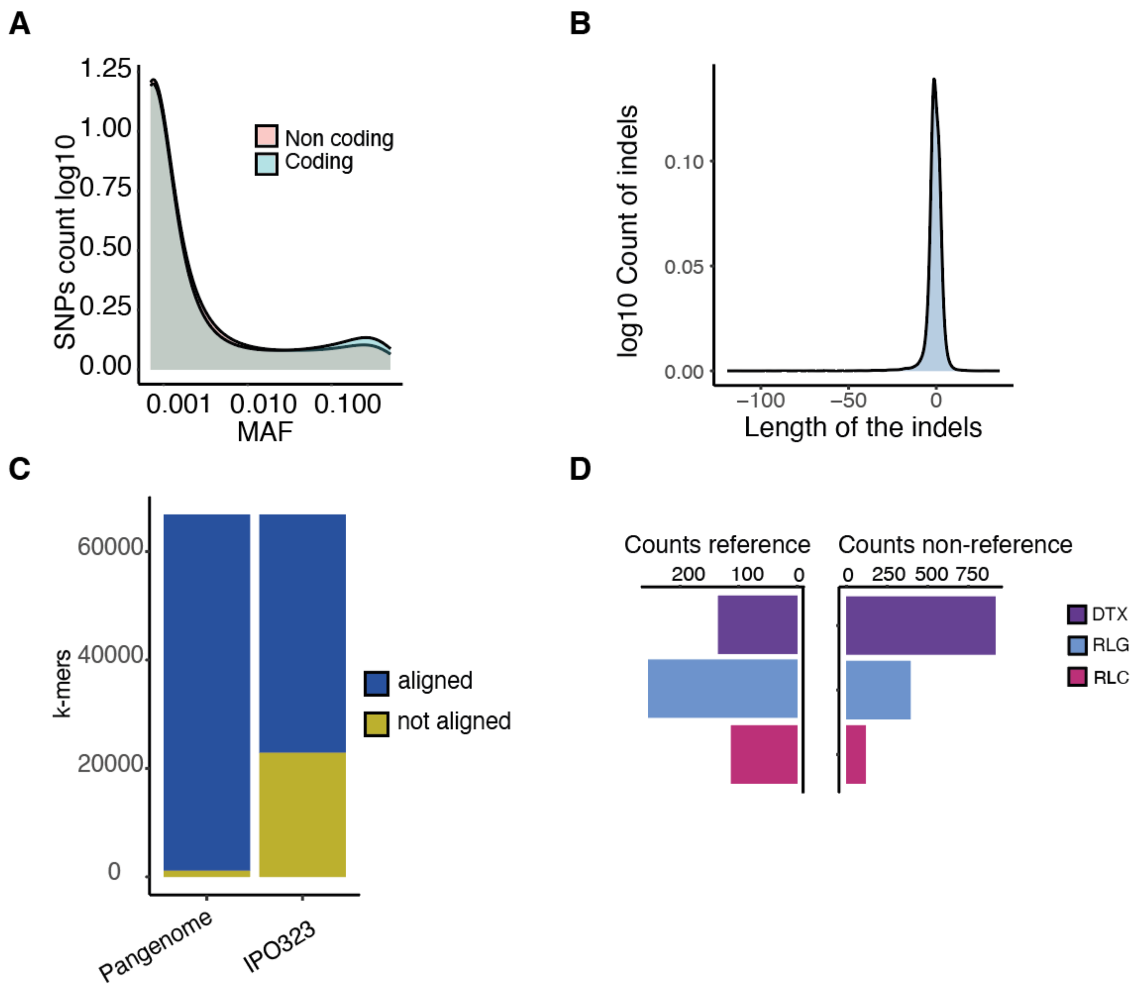


**Figure S2**: SNP and indel variant distribution in the European diversity panel. (A) Minor allele frequency distribution of coding and non-coding variants in the European diversity panel. (B) Indel length distribution. (C) Proportion of k-mers aligning (yellow) or lacking positioning (blue) on the reference genome (IPO323) and on the pangenome of the species. (D) Count of the most frequent TE superfamilies present in the European diversity panel split into reference and non-reference TEs.

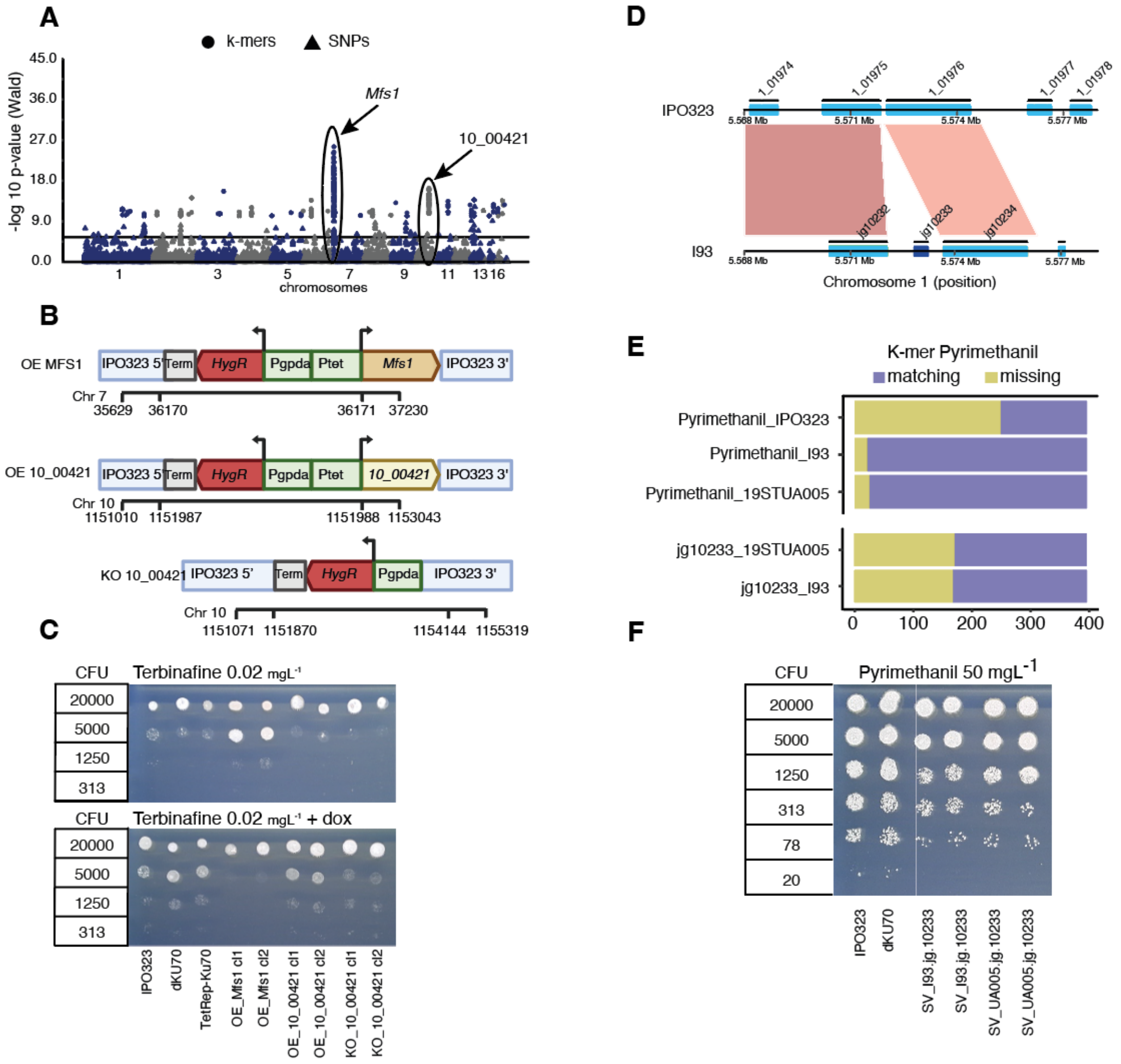


**Figure S3**: Profiling and functional validation of the role of the 10_00421 gene in terbinafine resistance. (A) Manhattan plot of significant SNPs (triangle) and k-mers (dots) for terbinafine relative growth values (0.1 mg.L$^{-1}$). Shapes identify variant types including SNPs (dots) and k-mers (triangle). (B) Construct design of OE_MFS1, OE_Zt09_10_00421, and KO_Zt09_10_00421 and (C) spotting assays on terbinafine (0.02 mg.L$^{-1}$), with and without doxorubicin (dox) (at 30 mg.L$^{-1}$). (D) Synteny plot of the chromosome 1 (5.568-5.577 Mb) of genomes IPO323 and I93. In light blue, gene regions; in dark blue is the gene jg.10233. (E) Proportion of k-mers aligning (yellow) or not (blue) to the genomes of IPO323, I93, and 19UA005; proportion k-mers aligning (yellow) or not (blue) to the structural variant (SV) affecting the gene jg.10233 in the strains I93 and 19UA005. (F) Spotting assay for pyrimethanil concentration (50 mg.L$^{-1}$).

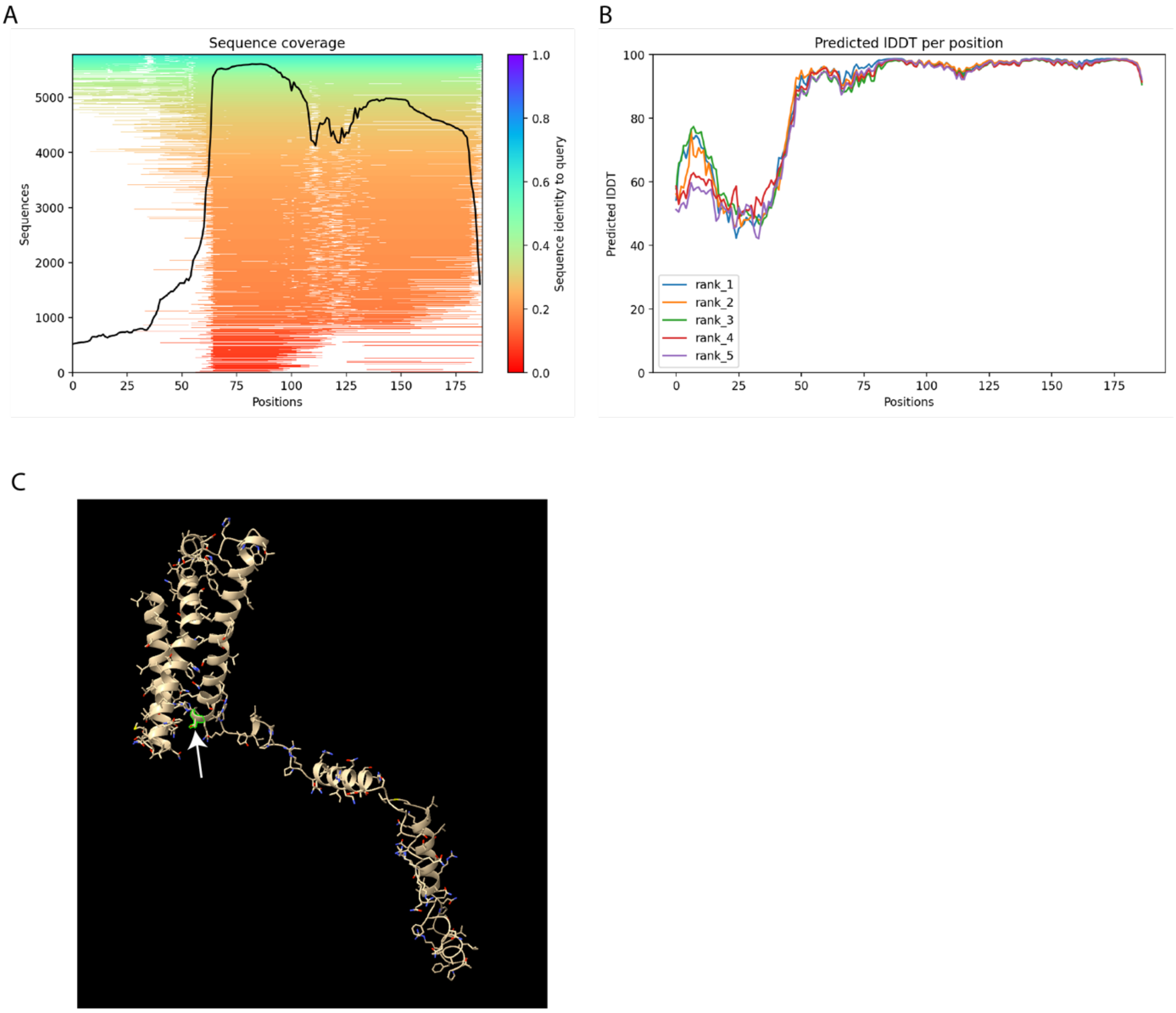


**Figure S4**: Predicted 3D structure of *sdhC1* generated with Alphafold2. (A) Number of sequences per position – Sequences are reliable with at least 30 sequences per position, for best performance, ideally 100 sequences. (B) Predicted lDDT per position - model confidence (out of 100) at each position. (C) The 3D structure of SDHC. Position 79 is highlighted in light green and at the beginning of the helical domain (white arrow).

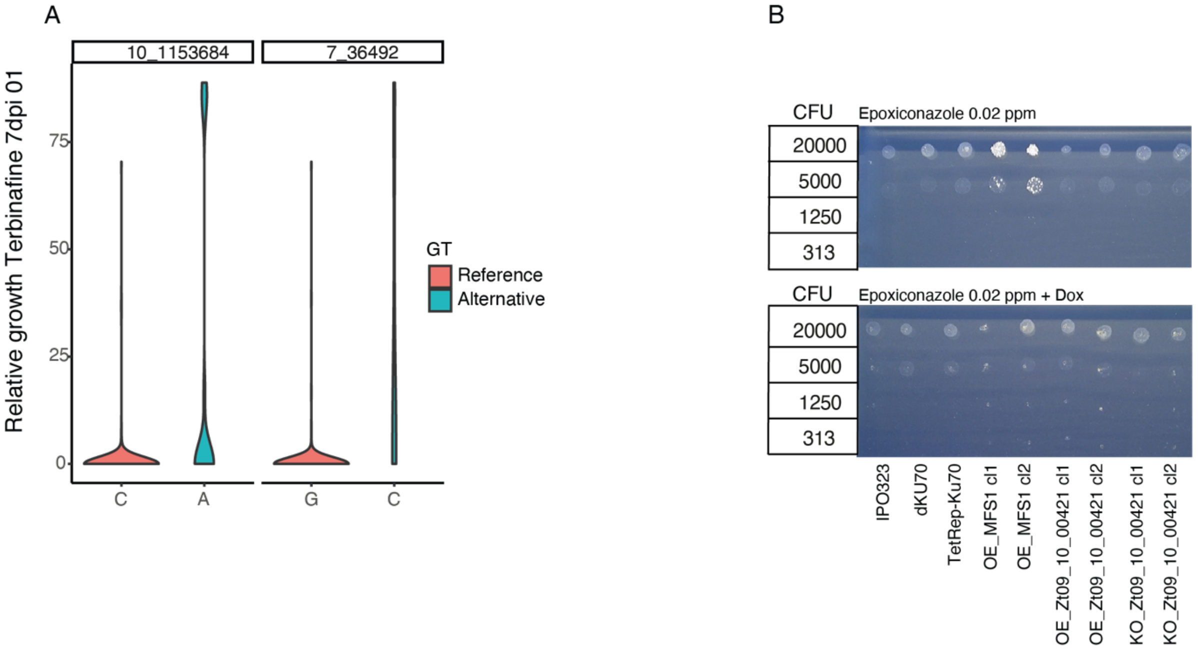


**Figure S5**: (A) Relative growth values of the most significantly associated SNPs in *Mfs1* (7_36492) and 10_00421 (10_1153684) in terbinafine (0.1 mg.L$^{-1}$). (B) Spotting assay for epoxiconazole resistance. The overexpression lines (OE) carry a Tet-Off, doxycycline-repressible promoter. The mutants tested are KO_Zt09_10_00421, OE_Zt09_10_00421, OE_MFS1, TetRep-Ku70, dKU70 and IPO323.

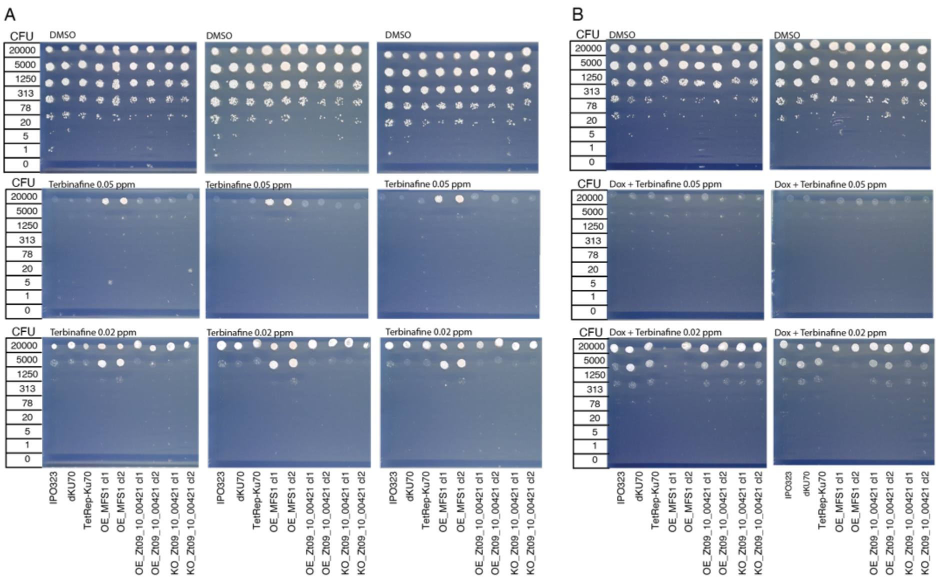


**Figure S6**: Spotting assay performed on AE media amended with (A) terbinafine (0.2 - 0.05 mg.L$^{-1}$) and (B) in presence of doxycycline (30 mg.L-1) with KO_Zt09_10_00421, OE_Zt09_10_00421, OE_MFS1, TetRep-Ku70, dKU70 mutants and IPO323.

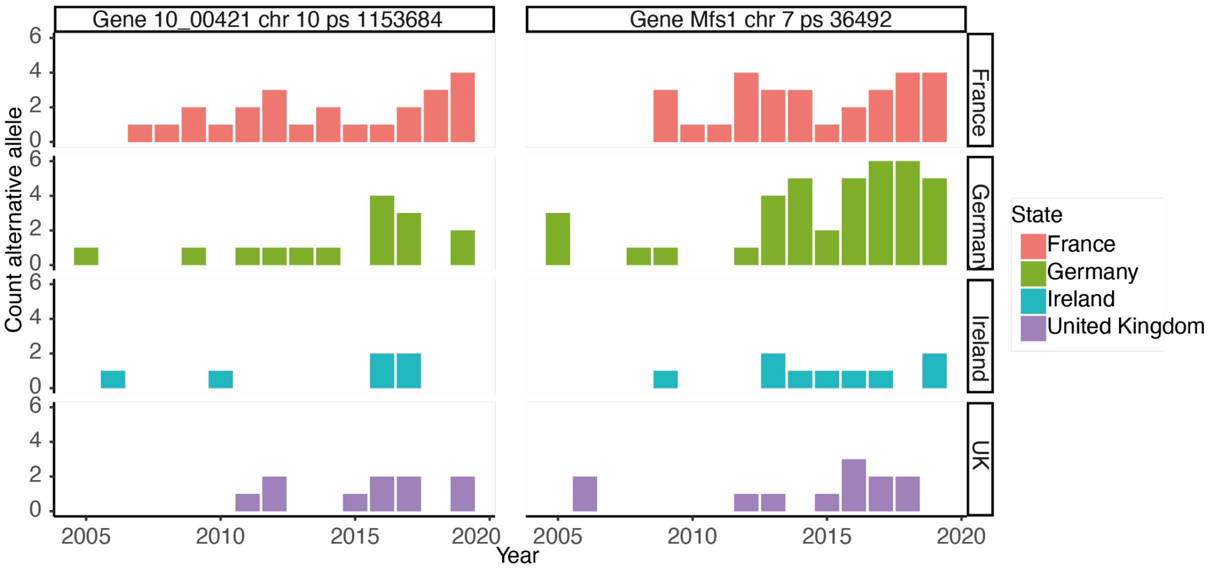


**Figure S7**: (A) Frequency of the most resistant alleles found for *Mfs1* (chr 7, position 36,492) and Zt_10_00421 (chr 10, position 1,153,684) in populations from France, UK, Ireland and Germany.

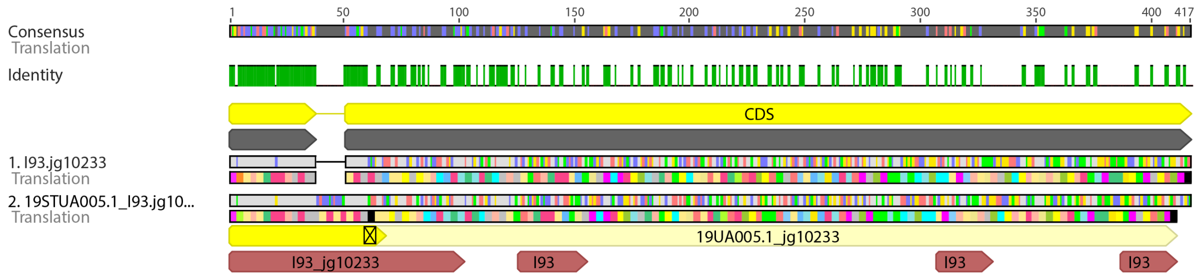


**Figure S8**: (A) Multiple sequence alignment of the genes jg10233 in reference genome isolate I93 and in 19STUA005.1. In yellow is the haplotype for jg10233 as discovered in isolate 19STUA005.1 and in brown the haplotype of jg10233 in the isolate I93. The black asterisk marks the presence of a stop codon.

### Supplementary table legends

**Table S1**: Overview of all *Z. tritici* strains used in this study reporting "Isolate_names_present_in_the_VCF": a unique ID for each strain. The columns "State", "Continent", "latitude" and "longitude" provides geographic information of each of the strains while the column "year" correspond to the year the strain was collected. The column "Published" and "Bioproject" provide further information to identify each of the strains used in this work. The columns reporting phenotyping information consist of the name of the fungicides, followed by the concentration, with the acronym C and followed by the days post inoculation (dpi) for the phenotyping. The columns "Strains_present_in_the_TE_GWAS", "Strains_present_in_the_Indels_GWAS", "Strains_present_in_the_CNV_GWAS", "Strains_present_in_the_SNPs_GWAS" and "Strains_present_in_the_kmer_GWAS" report strains included in different association studies.

**Table S2**: Overview of all fungicides tested. The column "Fungicide_ID" is matching with the fungicide's headers in Table S1. The column "common name" includes the fungicide name, followed by the time point (7-14dpi), concentration (mg.L$^{-1}$), name of the fungicides tested, days post inoculation (dpi), fungicide concentration, fungicide group name, matches to the function/chemical according to the FRAC guidelines and for the fungicides not present in the FRAC accordingly to the nomenclature accessible online. Pesticide use in the EU (2005-2019) information on whether each fungicide was applied and the respective information of the use in the EU (2005-2019; Eurostat data). Information on whether the fungicides were used in wheat fields and citations. Reported fungicide resistance in *Z.tritici* and citations.

**Table S3**: Frequency of indels in the diversity panel.

**Table S4**: Results of association mapping for significant k-mers above the 5% threshold across all tested fungicides and aligning to a unique position in the IPO323 reference genome using Bowtie2 v.2.3.3 ("-a --best –strata") (Langmead and Salzberg 2012). Further information includes the general output of Gemma (Zhou and Stephens 2012; Voichek and Weigel 2020), the 25-bp k-mer, the gene id of intersecting genes, and the fungicide_ID. Finally, the k-mer strand and the mismatch, corresponding to the position and base pairs swap compared to the reference.

**Table S5**: Significant k-mers aligning to the IPO323 reference genome or the full pangenome combined.

**Table S6**: Results of association mapping for significant TEs above the Bonferroni threshold for all tested fungicides. General output of Gemma (Zhou and Stephens 2012), including chromosome, position, major and minor alleles, p_score and other statistics. Further information includes fungicide_ID, transposable element (TE) identifiers, reference versus non-reference TE according to *ngs-te-mapper2* (Linheiro and Bergman 2012). A unique key for the TE identification including the TE superfamily, a unique ID , and the start/end coordinates are provided.

**Table S7**: Results of association mapping for significant CNVs above the Bonferroni threshold for all tested fungicides. General output of Gemma (Zhou and Stephens 2012), including chromosome, position, major and minor alleles, p_score, and other statistics. Further information includes the fungicide ID, intersecting genes and reference identifier in the IPO323 genome (Grandaubert, Bhattacharyya, and Stukenbrock 2015).

**Table S8**: Results of association mapping for significant SNPs above the Bonferroni threshold for all tested fungicides. General output of Gemma (Zhou and Stephens 2012), including chromosome, position, major and minor alleles, p_score, and other statistics. Further information includes the putative impact of each variant, as obtained by snpEff.

**Table S9**: Proportion of significant k-mers per fungicide mapping to each of the 19 reference genomes of the species pangenome (Badet et al. 2020). Generated with bowtie v1.2.2, with the command "-a –best –strata" (Langmead and Salzberg 2012).

**Table S10**: Proportion of significant k-mers for pyrimethanil resistance phenotyping and aligning to the reference genomes of IPO323, I93 or UA005, as well as matching the structural variant (SV) detected in I93 and UA005.

**Table S11**: Culture media used in this study and formulation.

**Table S12**: Correlation of colony growth between two independent replicates for 95 strains at 7 dpi. "Position" indicates the plate coordinates of each colony, while "test1" and "test2" report the measured colony areas in replicates 1 and 2, respectively.

**Table S13**: Summary of the number of synonymous and missense mutations generated by snpEff.

**Table S14**: Overview of all fungicides tested and a list of the association files. The column "Fungicide_ID " is matching with the headers in the Table S1. SNPs_GWAS_table is the list of Gemma association tables obtained using the SNP genotyping VCF file split per fungicide, concentration, and time point combination. Identical file sets were produced for Gemma association mapping tables under the columns Indel_GWAS, TE_GWAS, and CNV_GWAS. All_unique_K-mers_fasta, list of files containing 25-bp k-mers above the 5% threshold per fungicide. Pass_threshold_5_K-mers_GWAS, the list of output files of the K-mers above the 5% threshold per fungicide lacking any reference genome alignment. Match_K-mer_GWAS_unique, list of files containing the K-mers above the 5% threshold per fungicide, which align uniquely to the reference genome. Pangenome results, list of K-mers above the 5% threshold per fungicide, which align uniquely to the reference genome.

**Table S15**: Associations above the Bonferroni threshold per random subsample of the European diversity panel based on mefentrifluconazole resistance phenotyping. Output of Gemma (Zhou and Stephens 2012). Further information includes sample size (ranging from 100 to 800) and resampling set (1-100).

**Table S16**: Associations above the Bonferroni's threshold per random subsample of the European diversity panel based on prothioconazole resistance phenotyping. Output from Gemma (Zhou and Stephens 2012). Further information includes sample size (ranging from 100 to 800) and resampling set (1-100).

**Table S17**: Results of association mapping for significant k-mers above the 5% threshold for all fungicides. Further information include general output of Gemma (Zhou and Stephens 2012; Voichek and Weigel 2020).

**Table S18**: Results of association mapping for significant indels above the Bonferroni threshold and all fungicides tested. General output of Gemma (Zhou and Stephens 2012), including chromosome, position, major and minor alleles, p_score and standard Gemma statistics. Further information includes the fungicide IDs, intersecting genes, and the reference indels in the IPO323 reference genome (Grandaubert, Bhattacharyya, and Stukenbrock 2015).

**Table S19**: Mutant and plasmid identifiers used in this study.

**Table S20**: Primers used to check for homologous recombination of OE_MFS1, SDHC_Thr79Asn, OE_Zt09_10_00421, KO_Zt09_10_00421, BetaTUBGlu198Ala, SV_I93.jg.10233, SV_UA005.jg.10233 in the IPO323 background. "Position" indicates the location of the oligo on the target sequence while "OLIGO" corresponds to the name of the primer. "Len" is the length of the oligonucleotide (in bp), and "Tm" is the melting temperature (°C) of the oligo, indicating the temperature at which half of the oligo is bound to its complementary strand. "GC"% is the percentage of guanine (G) and cytosine (C) bases in the oligo, which influences stability and Tm. The column "Any_th" refers to the potential binding of the oligo to non-target sequences. The column "3_th", corresponds to the specificity at the 3' end of the oligo, which is crucial for proper binding and amplification. The column "Hairpin" indicates the presence or absence of hairpin structures in the oligo. Finally, "Seq" is the nucleotide sequence of the oligo. Further information includes the PCR polymerase used, either Q5 (Q5® High-Fidelity DNA Polymerase - New England Biolabs) or G2 (GoTaq® G2 Flexi DNA Polymerase - Promega).
